# Protective efficacy of mutant strains of *Borrelia burgdorferi* as potential reservoir host-targeted biologics against Lyme disease

**DOI:** 10.1101/2025.07.09.663885

**Authors:** Venkatesh Kumaresan, Trevor C. Smith, Miranda Lumbreras, Taylor MacMackin Ingle, Nathan Kilgore, Jolie F. Starling-Lin, Elizabeth J. Horn, J. Seshu

**Author notes:** Corresponding Author: J. Seshu, Department of Molecular Microbiology and Immunology BSE3.230; 1 UTSA Circle, The University of Texas at San Antonio San Antonio, TX-78249. Department of Molecular Genetics and Microbiology, Tufts University School of Medicine (Trevor C. Smith). Department of Immunology, Duke University Medical Center (Miranda Lumbreras).

## Abstract

Lyme disease (LD), caused by *Borrelia burgdorferi* (*Bb*), is the most common vector-borne disease in the United States. Novel strategies to control transmission of *Bb* to humans via ticks are critical for prevention of LD. One such strategy is to leverage non-infectious mutant strains of *Bb* for pathogen-derived biologics to block *Bb* transmission during the natural infectious cycle. Deletion of <u>B</u>orrelia host-<u>ad</u>aptation <u>R</u>egulator (Δ*badR*) and replacement of 8 conserved residues with alanines in Carbon Storage Regulator A of *Bb* (8S) resulted in mutants, with upregulation of RpoS and several immunogenic lipoproteins, that were incapable of colonization in C3H/HeN mice. Intradermal vaccination of mice with the live mutant strains resulted in significant anti-borrelial antibody responses and a reduction in *Bb* acquisition by naïve *Ixodes scapularis* larvae following needle challenge with *Bb* B31-A3. While vaccination with Δ*badR* mutant conferred significant reduction in the percentage of infected mice, ML23 and 8S mutants conferred variable levels of protection. Comparative proteomic analysis of Purified Borrelial Lipoproteins (PBLs) from parental and mutant strains revealed similar and unique antigenic components. Infection derived mouse serum and serum from Lyme disease patients exhibited reactivity to lysates and PBLs from these mutants. Immunization with PBLs from B31-A3 and 8S strains conferred significant protection against challenge with *Bb* infected nymphs, underscoring the utility of PBLs as protective formulations. Overall, non-infectious mutant strains, or their lipoproteins, can be exploited as biologics to block the acquisition of *Bb* by naïve larvae from reservoir hosts, disrupting the enzootic transmission cycle of the agent of Lyme disease.

## Introduction

Lyme disease is the most common vector-borne disease in the US with 89,000 cases reported to Centers for Disease Control and Prevention (CDC, Atlanta) in 2023. It is estimated that around 476,000 cases occur each year, reflecting an increase in the incidence of Lyme disease (1, 2). *Borrelia burgdorferi* (*Bb*), the spirochetal agent of Lyme disease, is transmitted to vertebrate hosts through the bite of infected *Ixodes scapularis* (*Is*) ticks, which are known to transmit up to 7 pathogens to humans and naïve vertebrate hosts (3). Currently, there are no vaccines available for use in humans. Hence, there is a dire need to advance strategies to limit or completely interfere with the transmission of one or more tick-borne pathogens as they cycle through *Is* ticks and a variety of vertebrate hosts (4, 5). One strategy to combat Lyme disease is to reduce the burden of spirochetes in reservoir hosts in endemic areas, with the expectation that reduced transmission via ticks will gradually reduce the incidence in humans (6, 7). Amongst a vast number of mutant strains of *Bb* that have been generated thus far, several are incapable of survival in mammalian hosts or have a highly attenuated phenotype (8–10). However, these mutant strains, or their subcellular products, provide a rich assortment of candidates that can be tested for efficacy as reservoir host-targeted biologics, exploiting their inability to colonize mammalian hosts while capable of inducing protective immune responses.

Mutant strains of *Bb* have been utilized to dissect key features of innate and adaptive immune responses, as well as metabolic fitness to survive during the tick and mammalian phases of infection. Prior to establishment of tools and methodologies for targeted manipulation of the genome of *Bb*, a non-motile, flagella-less mutant was isolated and evaluated for protective efficacy as a live-attenuated vaccine in murine models of Lyme disease (11, 12). Additional mutant strains lacking one or more plasmids encoding for major lipoproteins of *Bb* such as <u>O</u>uter <u>s</u>urface <u>P</u>rotein <u>A</u> and <u>B</u> (OspA; OspB) were evaluated for their protective capabilities (13–15). Moreover, an inactivated, live reservoir host-targeted whole cell vaccine based on heterologous expression of OspA as a recombinant protein in *E. coli* has also been shown to significantly reduce the levels of *Bb* positive *Is* larvae parasitizing *Peromyscus leucopus (P. leucopus)* hosts (16). Mutant strains of *Bb* either lacking *p66* or with hyperexpression of *p66* that were incapable of establishing infection in a murine model of Lyme disease, conferred protection following multiple immunizations in C3H/HeN mice against challenge with the parental strain (17). The rapid clearance of the *p66* mutant from the skin of C3H/HeN mice was not influenced by the skin antimicrobial peptide mCRAMP and was unable to colonize neutrophil depleted mice, suggesting a novel role for *p66* in the colonization of *Bb* (18). Furthermore, another borrelial mutant lacking expression of abundant major surface-exposed lipoproteins (OspA, OspB and OspC, *ospABC^-^* mutant) was used to identify non-abundant immunogenic proteins, expanding the utility of mutant strains of *Bb* in developing tools or preparations that can be used to induce protective immune responses (19). Two less abundant proteins identified from the above study, BBA34 and BB0238, were shown to be immunogenic during the course of mammalian infection (20, 21) These studies provide a basis for evaluating the utility of additional borrelial mutants to explore the host responses induced due to the lack or overexpression of select determinants and to examine the ability of these mutants, or their subcellular components, to serve as pathogen-derived biologics capable of altering host responses to limit colonization of wild type *B. burgdorferi*.

Previously it was shown that the deletion of <u>B</u>orrelia host-<u>ad</u>aptation <u>R</u>egulator (Δ*badR* mutant) results in de-repression of RpoS and an increase in the levels of expression of several major borrelial lipoproteins that are known to play a critical role in mediating the interactions of *Bb* with host cell surfaces (22, 23). Levels of <u>O</u>uter <u>s</u>urface <u>p</u>rotein <u>C</u> (OspC), Decorin <u>b</u>inding <u>p</u>roteins <u>A</u> and <u>B</u> (DbpA/B) and BBK32, among others, were all upregulated in the *badR* mutant. In another mutant, substitution of eight conserved residues with alanines in Carbon Storage Regulator A of *B. burgdorferi* (CsrA*_Bb_*; 8S mutant) also resulted in increased levels of expression of major lipoproteins of *Bb* (24, 25). Both these mutant strains are incapable of colonization of mouse models of infection and are therefore attenuated for infection, likely due to either a metabolic defect in the mutant for survival within mammalian hosts or to innate/adaptive immune mechanisms clearing the spirochetes that hyper-express immunogenic borrelial lipoproteins and other immunologically relevant determinants (22, 24). However, the increased levels of several RpoS-regulated borrelial lipoproteins in both *badR* and 8S mutants could be exploited as biologics. Since surface-exposed borrelial lipoproteins elicit robust antibody responses in mammalian hosts, they are likely to confer protection against challenge with infectious *Bb* strains. In addition to the above mutants, a clonal isolate of *Bb* strain B31 lacking the 25kb linear plasmid (lp25, ML23) or an lp25-deficient clone from *Bb* strain B31-A3, which is incapable of colonization of the mammalian host, was also used in the study (26, 27). The lack of colonization of ML23 in C3H/HeN or the SCID mice is due to the deficiency of a key enzyme nicotinamidase (encoded by *bbe22* or *pncA* on lp25) critical for metabolism of nicotinamide to generate NAD (26–28).

Several studies have demonstrated the relevance of the antibody responses directed at surface-exposed borrelial lipoproteins in preventing transmission of *Bb* from ticks (29, 30) Therefore, this study is directed at exploiting borrelial mutants that hyper-express a large number of immunogenic borrelial lipoproteins as a biologic to prevent or reduce the pathogen burden and limit the ability of naïve larvae to acquire *Bb* from reservoir hosts. While several RpoS-induced lipoproteins were differentially expressed in both *badR* and *8S* mutants, the levels of <u>O</u>uter <u>s</u>urface <u>p</u>rotein <u>A</u> (OspA), a major immunogenic lipoprotein, was not altered in these mutants in comparison to the parental wild-type strain under *in vitro* growth conditions (22). Therefore, OspA-specific protective responses, essential for blocking *Bb* transmission from ticks to hosts, are retained in the mutants and may confer protection against *Bb* following intradermal vaccination (31–33). Moreover, a genome-wide proteomic analysis of *Bb* (B31) using sera from patients with natural infections revealed approximately 15% of the 1,292 evaluated open reading frames code for immunogenic products (34, 35). The *Bb* genome encodes more than 120 lipoproteins, constituting nearly 8% of open reading frames. Experimental examination of 125 lipoproteins revealed that 86 are secreted to the bacterial surface (34). Consequently, lipoproteins are critical antigenic components of *Bb*, demonstrating efficacy in inducing protective immunity both as individual antigens and as a cocktail of recombinant lipoproteins (36). Therefore, we hypothesized that these mutant strains, incapable of survival in mouse models of Lyme disease despite hyper-expression of major surface exposed lipoproteins, likely serve as potential biologics to interfere with pathogen survival and transmission from reservoir hosts (7, 30, 31, 37). Moreover, numerous lipoproteins with native composition and distribution/stoichiometry on borrelial outer membranes are expected to induce robust protective immunity (38). Exploring the protective efficacy of lipoproteins purified from borrelial mutants holds significance, providing valuable insights into the potential use of these strains as intact spirochetes or utilizing purified lipoproteins derived from these mutants as reservoir host-targeted vaccines.

In this study, we tested the hypothesis that vaccination with mutant strains of *Bb* that hyper-express a large number of lipoproteins stimulate protective immune responses that block the acquisition of *Bb* by naïve *Is* larvae. We assessed the utility of these mutant strains in providing protection against *Bb* infection in C3H/HeN and in transmission of *Bb* to naïve larvae following intradermal injection of intact spirochetes. We also determined the antigenicity of purified lipoproteins from wildtype (B31-A3), 8S and Δ*badR* strains using infection derived mouse serum and bar-coded Lyme disease patient serum (blinded with no patient information). Furthermore, the protective efficacy of the lipoproteins purified from these mutant strains following immunization via the intradermal route was determined against *Bb* infected tick challenge. Correlates of the immune response from these studies will aid in formulating purified borrelial lipoproteins for delivery via the oral route and in the validation of efficacy of peripheral immune responses induced against *Bb.* These studies are aimed at advancing methods to reduce pathogen burden in reservoir hosts, thereby reducing transmission of *Bb* to humans and naive reservoir hosts via ticks.

## Results

### Protective efficacy of mutant borrelial strains

We hypothesized that 8S and Δ*badR* strains can be exploited as live attenuated strains to induce protective antibody responses, a key correlate of protection, in C3H/HeN following two immunizations, as both these strains were incapable of survival in murine hosts. The ability of immunized animals to prevent acquisition of *Bb* by naïve larvae was used to measure the efficacy of the mutant strains to confer protective responses (Fig. 1A). As a control, a clonal isolate ML23 lacking lp25 but has lp28-1 from *Bb* strain B31 (26, 27). Two immunizations with 1×10^5^ *Bb*/mouse at day 0 and 16 resulted in significant levels of total IgG against B31-A3 lysate at 28 days post-vaccination (dpv) compared to unvaccinated controls (Fig. 1B). Antibody titers remained high in vaccinated mice at 42 dpv (data not shown). Mice were needle-challenged with 1×10^5^ *Bb* strain B31-A3 spirochetes per mouse on day 56 and *Is* larvae placed on mice on day 84 fed to repletion and genomic DNA isolated from four replete larvae from each mouse was used to quantify *Bb* burden via qPCR (Fig. 1C). Both unvaccinated mice allowed naïve larvae to acquire *Bb*, while only one out of three mice vaccinated with either 8S or Δ*badR* had fed larvae positive for *Bb.* All three mice vaccinated with ML23 prevented acquisition of *Bb* by naïve larvae (Fig. 1C). The skin samples from mice immunized with ML23 had no detectable *Bb*-specific DNA, while two mice were positive from those immunized with 8S or Δ*badR* even though there were no detectable *Bb* DNA in ticks fed on one of these mice (Fig. 1D). Moreover, *Bb*-specific DNA was detected in the lymph node and joints of one mouse vaccinated with 8S or Δ*badR,* respectively, consistent with the presence of *Bb* in fed larvae from these mice, while all three mice vaccinated with ML23 were negative (Fig. 1E and F). It was possible to consistently detect *Bb*-specific DNA in both larvae fed on unvaccinated mice as well as from different tissues (Fig. 1C, D, E, and F). These observations established the levels of protective immune responses induced following immunization with mutant strains of *Bb* that are incapable of colonization within murine hosts.

**Figure 1:**
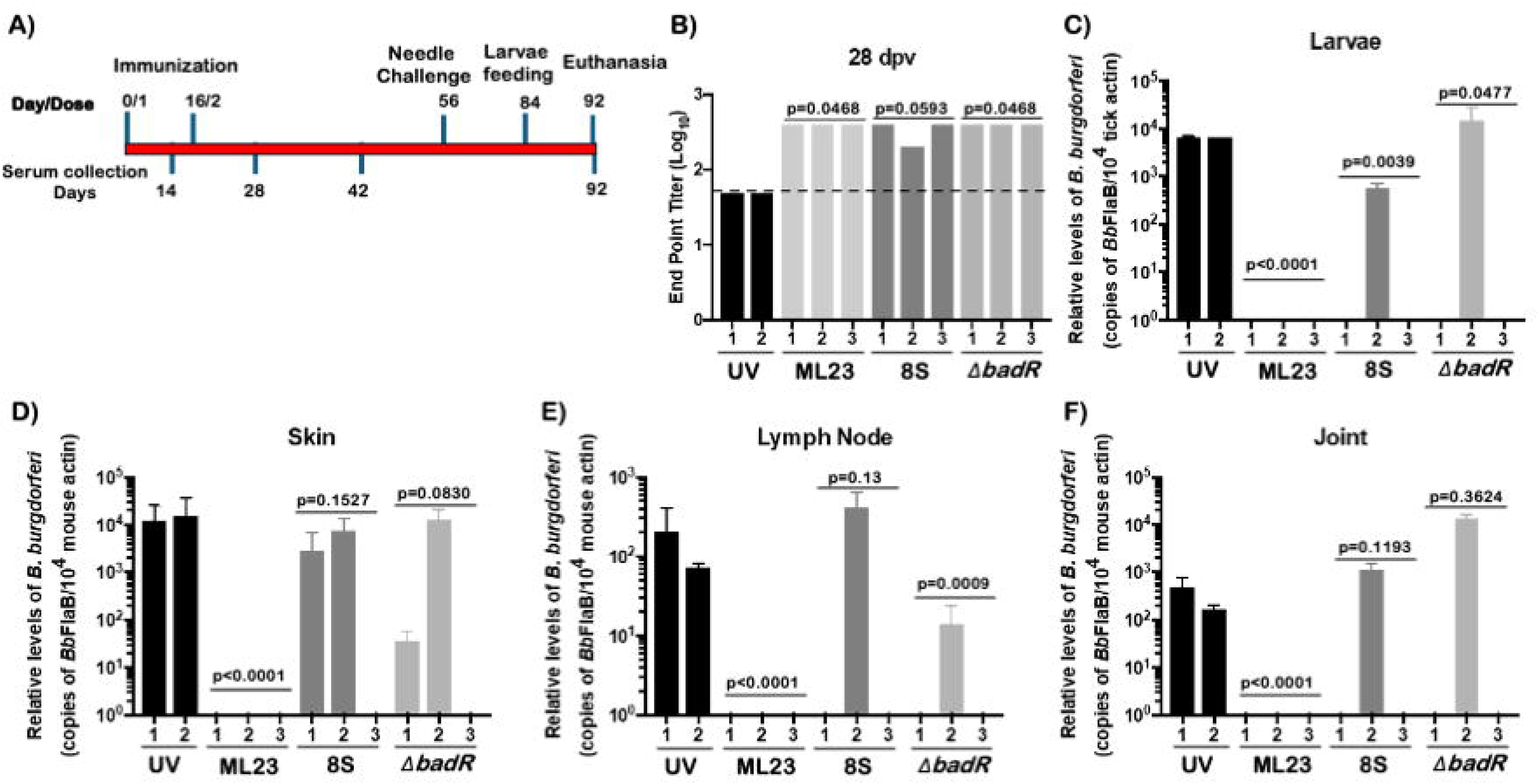
Levels of protection conferred by live attenuated strains in C3H/HeN mice against challenge (4 weeks) with *Bb* strain B31-A3. 1A) Graphical representation of the timeline of vaccination, needle challenge, larvae feeding and time of euthanasia of C3H/HeN mice. Serum collection timeline is shown at the bottom. 1B) Bar graphs showing the end-point titers of antibodies induced in immunized mice at 28 days post first immunization (dpi) determine using ELISA with *Bb* strain B31-A3 lysate as antigens. Differences between vaccinated and unvaccinated sample groups were analyzed using a Mann-Whitney U test; corresponding *p* values are shown at the bottom of each graph. A *p* value < 0.05 was considered statistically significant. 1C) Bar graphs showing *Bb* burden in *I. scapularis* larvae fed on challenged mice at 84 days post first immunization. Numbers of borrelial *flaB* copies were normalized against 10^4^ total tick actin copies. 1D) Bar graphs showing *Bb* burden in mouse tissues at the time of euthanasia. Numbers of borrelial *flaB* copies were normalized against 10^4^ total mouse actin copies in skin, lymph nodes and joints. For tissue and larval genomic DNA analyses, differences between vaccinated and unvaccinated groups were evaluated using a two-tailed Student’s *t*-test. A *p* value < 0.05 was considered statistically significant.

In the second experiment, mice were vaccinated at day 0 and 18 and needle challenged with 1×10^5^ infectious B31-A3 at day 92, followed by tick feeding at day 120 to determine the levels of protection over a longer duration of time (Fig. 2A). As expected, there were higher antibody titers in vaccinated animals at day 28 (Fig. 2B). There was a significant reduction in the *Bb* burden (p<0.05) by qPCR in larvae fed on vaccinated mice immunized with ML23, 8S or Δ*badR* compared to those fed on unvaccinated mice, suggesting that the duration of the immune response after the second immunization likely influenced *Bb* acquisition by naïve larvae (Fig. 2C). Although larvae fed on Δ*badR* vaccinated mice had detectable levels of *Bb*-specific DNA, qPCR analysis of skin and joint did not reveal presence of *Bb*-specific DNA at day 129 following euthanasia, although one mouse had detectable levels in the lymph node (Fig. 2D-F). However, *Bb*-specific DNA was detectable in several tissues from mice vaccinated with ML23 and 8S, reflecting increased levels of pathogen burden in these mice. The levels of significance were calculated based on comparison between vaccinated and unvaccinated groups using a two-sided Fisher’s exact test.

**Figure 2:**
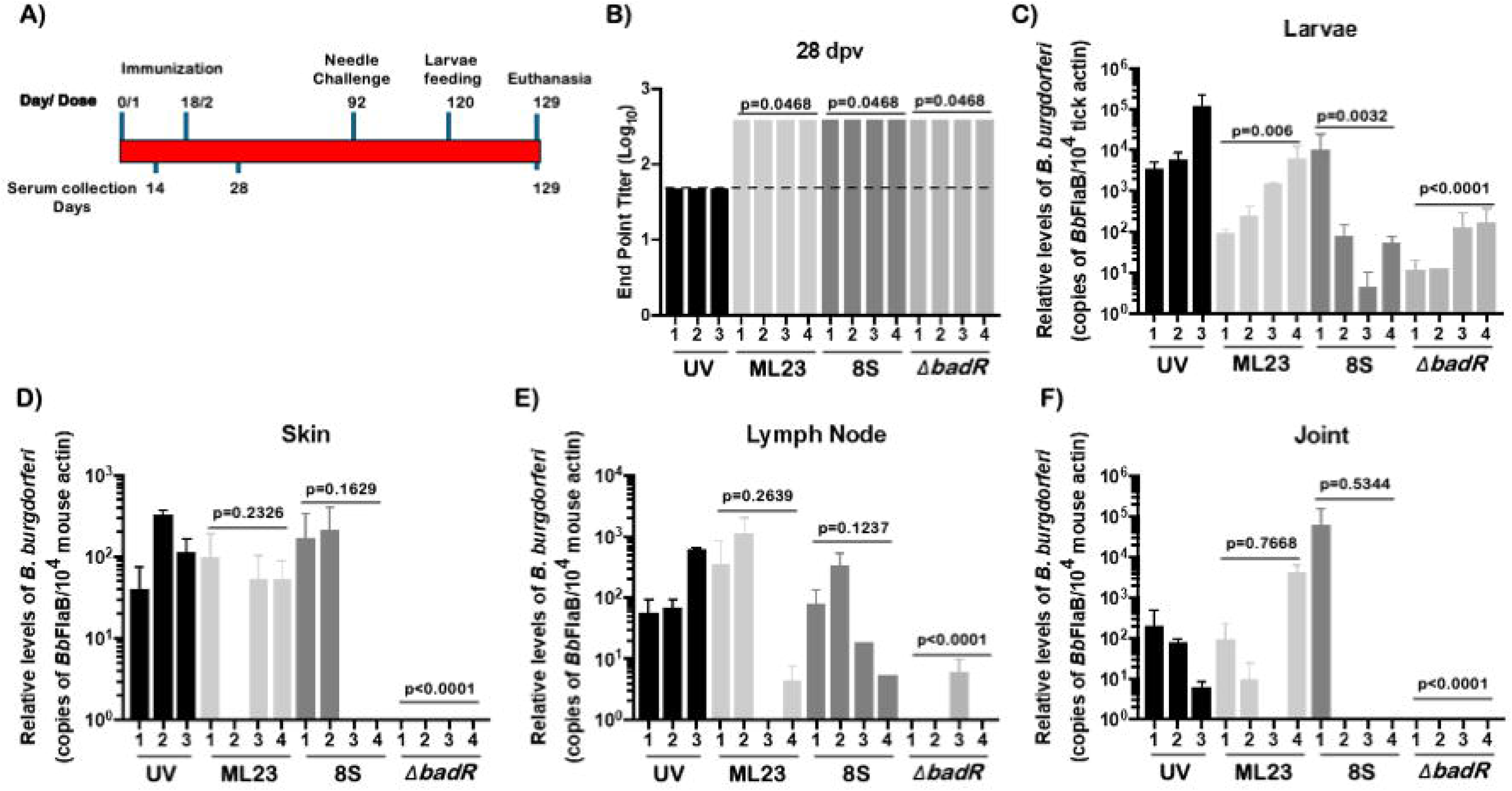
Levels of protection conferred by live attenuated strains in C3H/HeN mice against challenge (10 weeks) with *Bb* strain B31-A3. 2A) Graphical representation of the timeline of vaccination; needle challenge, larvae feeding and time of euthanasia of C3H/HeN mice. Serum collection timeline is shown at the bottom. 2B) Bar graphs showing the end-point titers of antibodies induced in immunized mice at 28 days post-first immunization (dpi) determined using ELISA with *Bb* strain B31-A3 lysate as antigen. Differences between vaccinated and unvaccinated sample groups were analyzed using a Mann-Whitney U test; corresponding *p* values are shown at the bottom of each graph. A *p* value < 0.05 was considered statistically significant. 2C) Bar graphs showing *Bb* burden in *I. scapularis* larvae fed on challenged mice at 84 dpi. Numbers of borrelial *flaB* copies were normalized against 10^4^ total tick actin copies. 2D) Bar graphs showing *Bb* burden in mouse tissues at the time of euthanasia. Numbers of borrelial *flaB* copies were normalized against 10^4^ total mouse actin copies in skin, lymph nodes and joints. For tissue and larval genomic DNA analyses, differences between vaccinated and unvaccinated groups were evaluated using a two-tailed Student’s *t*-test. A *p* value < 0.05 was considered statistically significant.

Tissue samples from the vaccinated/challenged mice were aseptically isolated following euthanasia at day 92 (Fig. 1A) or day 129 (Fig. 2A) and processed to recover viable spirochetes in BSK-II growth medium. Cultures were blindly passaged after 5 days into fresh BSK-II growth medium to minimize the toxicity associated with the degradation of host tissues and to facilitate growth of spirochetes. As shown in Table 1, 5 out of 5 unvaccinated mice challenged with *Bb* B31-A3 had viable spirochetes while only one mouse vaccinated with Δ*badR* was culture positive for spirochetes. Mice vaccinated with 8S (3/7) and ML23 (4/7) had tissues positive for spirochetes, suggesting reduced levels of efficacy in curtailing *Bb* survival with a challenge dose of 1×10^5^ spirochetes of *Bb* strain B31-A3. Since the data presented in Table 1 is consolidated from two separate experiments, as detailed in Figures 1 and 2, mice with a single tissue positive for *Bb* were considered as being infected. Moreover, mice with culture positive tissues were the ones that allowed for *Bb* acquisition by naïve larvae and correlated with high *Bb* load in those tissues by qPCR. While Δ*badR* vaccinated mice had significantly lower levels of infection (p=0.01), there was no significant reduction in levels of infection as determined by viable spirochetes in different tissues following vaccination with ML23 or 8S, although there was a 43% and 57% decrease in levels of infection compared to unvaccinated control mice, respectively. These data sets revealed that immunization with mutant strains of *Bb* results in select correlates of protection that can be exploited for developing reservoir host-targeted biologicals for reducing *Bb* burden. Although 8S and Δ*badR* mutants induced protective immune responses against WT-*Bb*, live attenuated strains induced immune responses and correlates of protection that varied between experiments. We then pursued evaluation of the immunogenic subcellular components, namely lipoproteins, purified from these mutants as protective pathogen derived biologics to block *Bb* transmission from vertebrate hosts to ticks.

**Table 1:** Efficacy of protection conferred by mutant strains of *Bb* following needle challenge with *Bb* strain B31-A3^$^.

| # of cultures positive /total # |  |  |  |  |  |  |  | # of mice<br>infected<br>/total # of<br>mice | Percent<br>of<br>infected<br>mice |
| --- | --- | --- | --- | --- | --- | --- | --- | --- | --- |
|  | Skin | Lymph<br>node | Spleen | Bladder | Heart | Joint | All<br>Sites |  |  |
| Unvaccinated | 5/5 | 5/5 | 1/5 | 5/5 | 5/5 | 5/5 | 26/30 | 5/5 | 100 |
| ML23 | 4/7 | 4/7 | 3/7 | 4/7 | 4/7 | 4/7 | 23/42 | 4/7 | 57<br>(p=0.2) |
| 8S | 3/7 | 3/7 | 1/7 | 3/7 | 3/7 | 2/7 | 15/42 | 3/7 | 43<br>(p=0.08) |
| $\Delta badR$ | 1/7 | 1/7 | 0/7 | 1/7 | 1/7 | 1/7 | 5/42 | 1/7 | 14<br>(p=0.01) |

<sup>§</sup>Consolidated representation of spirochetal growth in BSKII medium following propagation of tissues isolated from C3H/HeN mice used for infectivity analysis described in Figures 5 and 6.

### Extraction and profiling of borrelial lipoproteins

Lipoproteins were extracted from *Bb* strain B31-A3 (wild type parental strain used to generate the mutants), 8S and Δ*badR* using Triton X-114, and the protein profiles were compared between the insoluble pellet, aqueous phase (water-soluble proteins), and TX-114 phase (detergent phase) as described earlier (39). The total protein profile of the partitioned proteins separated on an SDS-PAGE gel and stained with Coomassie blue were distinct among all three strains (Fig. 3A). Immunoblot analysis using a high dilution (1:10,000) of *Bb* strain B31-A3 infection-derived mouse serum from C3H/HeN mice revealed high reactivity with proteins in the detergent phase in all three strains (Fig. 3B). Moreover, immunoblot of these partitioned proteins revealed the presence of FlaB (major flagellin of *Bb*), predominantly in the pellet and aqueous phase fractions, with no apparent levels of FlaB detected in the detergent phase fraction (Fig. 3C). Since much of this study is focused on purified borrelial lipoproteins from *Bb*-B31, 8S and Δ*badR* strains, we compared the detergent phase proteins from these strains and noted apparent differences between them (Fig. 4A). The infection-derived serum had the most reactivity with the detergent phase proteins, as a majority of these proteins are surface exposed lipoproteins that are immunogenic. Notably, there are increased levels of an approximately 33 kDa protein that is present in the detergent fraction of 8S compared to wild type or Δ*badR* strains (Fig. 4A). Based on the size and mass spectrometric analysis, this protein was determined to be Outer surface protein B (OspB), which has been shown to be an immunogenic surface exposed lipoprotein impacting antibody dependent mechanisms limiting *Bb* survival (Fig. 4B). While OspB was apparent in detergent phase extracts of the 8S strain, both the WT and Δ*badR* strains expressed OspB as one of the five most abundant lipoproteins (Fig. 4C). Three independently purified TX-114 fractions from all three strains were analyzed by unbiased proteomics and showed similar protein abundance, with peptides derived from major surface exposed lipoproteins, OspA, B, C, D and LP6.6 being the most abundant, validating the detergent extraction procedure (Fig. 4D).

**Figure 3:**
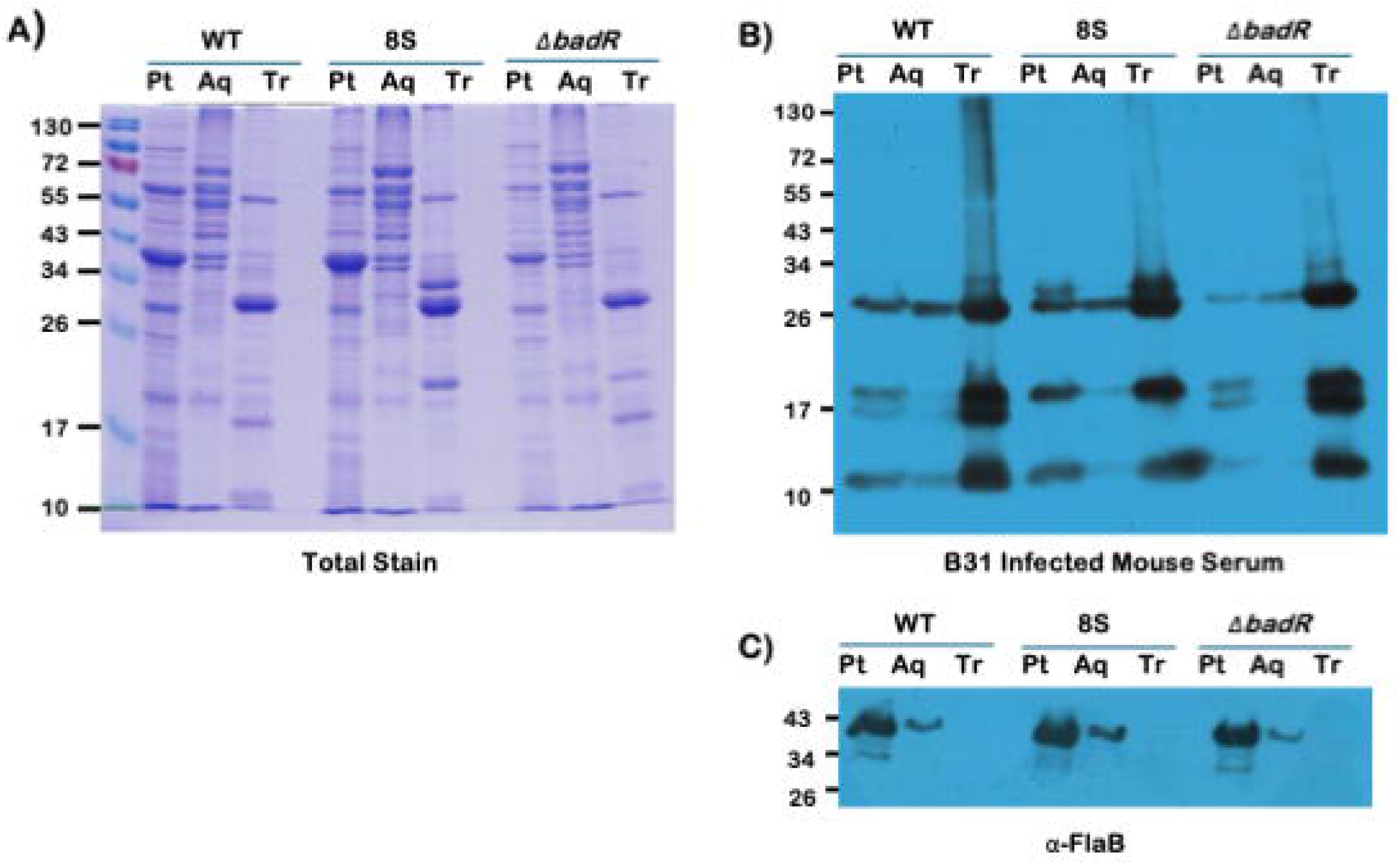
Extraction and Immunoreactivity of Borrelial Lipoproteins. *B. burgdorferi* (*Bb*) strains B31-A3 (WT), Δ*badR* and 8S were propagated at 32°C, pH7.6 to a density of 1×10^8^ *Bb*/mL cells. The cultures were washed with HBSS and partitioned into aqueous and detergent phase using Triton X-114 and were separated using SDS-12.5%PAGE gel. 3A) Total stain showing the proteins partitioned as the insoluble pellet (Pt), aqueous (Aq) and Triton X-114 (Tr) phases from all three strains. Immunoreactivity of partitioned proteins with 3B) B31-A3 infected mouse serum and 3C) anti-FlaB mouse serum. Molecular weights in kDA are indicated on the left.

**Figure 4:**
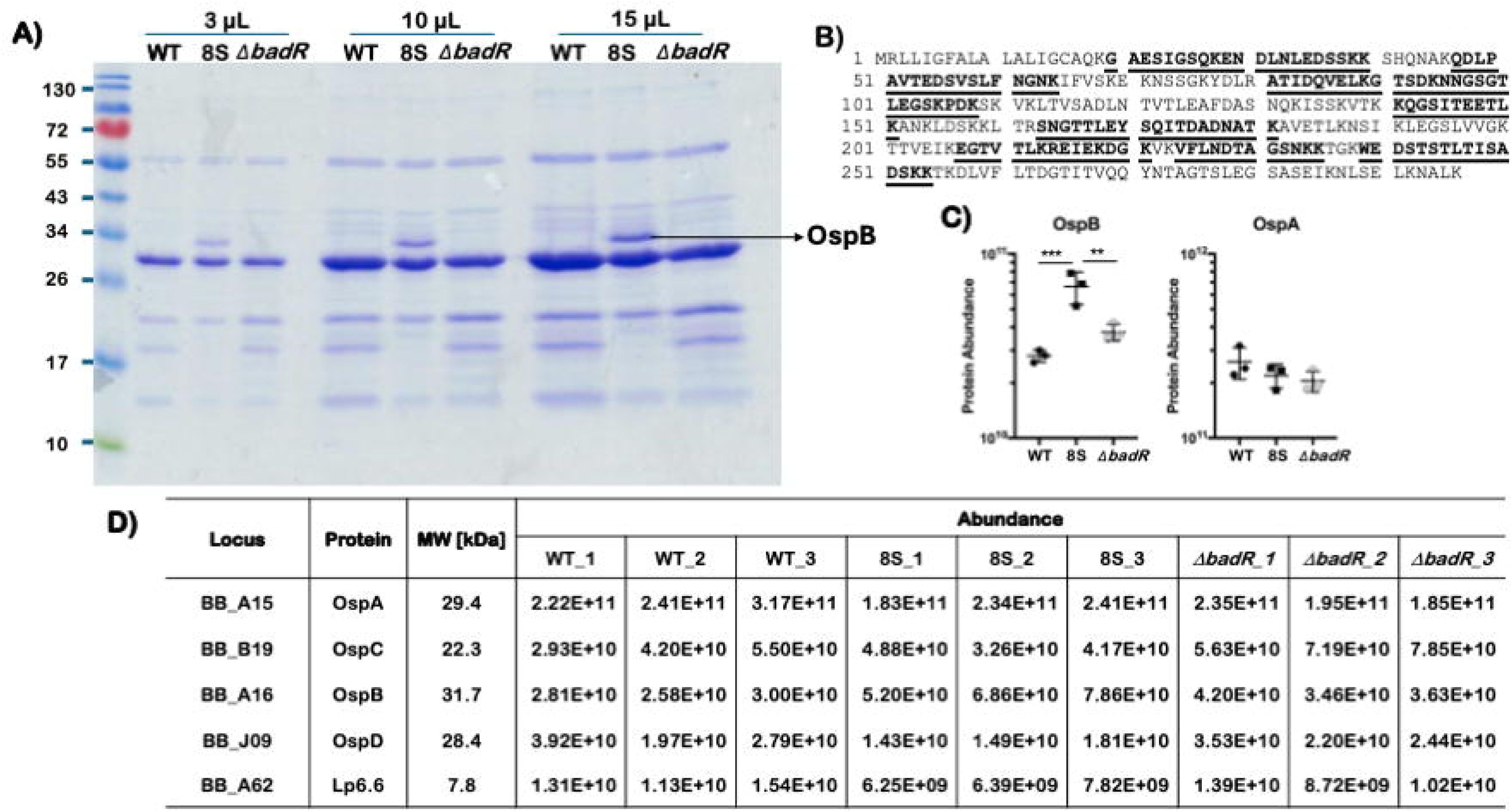
Purification and abundance of borrelial lipoproteins from mutant strains of *B. burgdorferi*. Total lysates of *B. burgdorferi* cultures were partitioned using Triton X-114 and the detergent phase proteins were subjected to proteomic analysis. 4A) Total stain of Triton X-114 phase proteins from WT (B31-A3), 8S and Δ*badR* separated using SDS-12.5% PAGE and stained with Coomassie Brilliant Blue. 4B) Mass spectrophotometric analysis of a detergent phase protein expressed at a higher level in the 8S mutant was identified as OspB. 4C) Relative abundance of OspB and OspA in WT, 8S and Δ*badR*. 4D) Peptide abundance reflecting the expression of 5 predominant lipoproteins in the detergent phase extract of borrelial mutants determined by proteomic analysis.

### Reactivity of serum from Lyme disease patients

Human serum samples with no patient identifiers or clinical information from patients with or without Lyme disease were obtained from the BioBank supported by the Bay Area Lyme Foundation and tested for reactivity with lipoproteins from parental and mutant strains using an ELISA-based method (Fig. 5). While several human serum samples showed high titers of IgG specific to lipoproteins from all three strains, there were several samples that had baseline reactivity similar to a normal human serum sample from a local blood bank used as a negative control (normal serum). There were differences in the end-point titers of human serum samples against lipoproteins, suggesting differences in either the specificity of the antibodies or potential variations in the antigenic profiles of the three strains. Human serum samples that had significantly higher titers (*p* value of <0.05) against lipoproteins from the parental wild-type strain at levels above base line titers had similar reactivity with lipoproteins from 8S or Δ*badR* strains compared to human serum samples that were classified as negative (Fig. 5). Immunoblot analysis of a positive human serum sample (LD585) showed reactivity to lipoproteins of all three borrelial strains (Fig. 5B).

**Figure 5:**
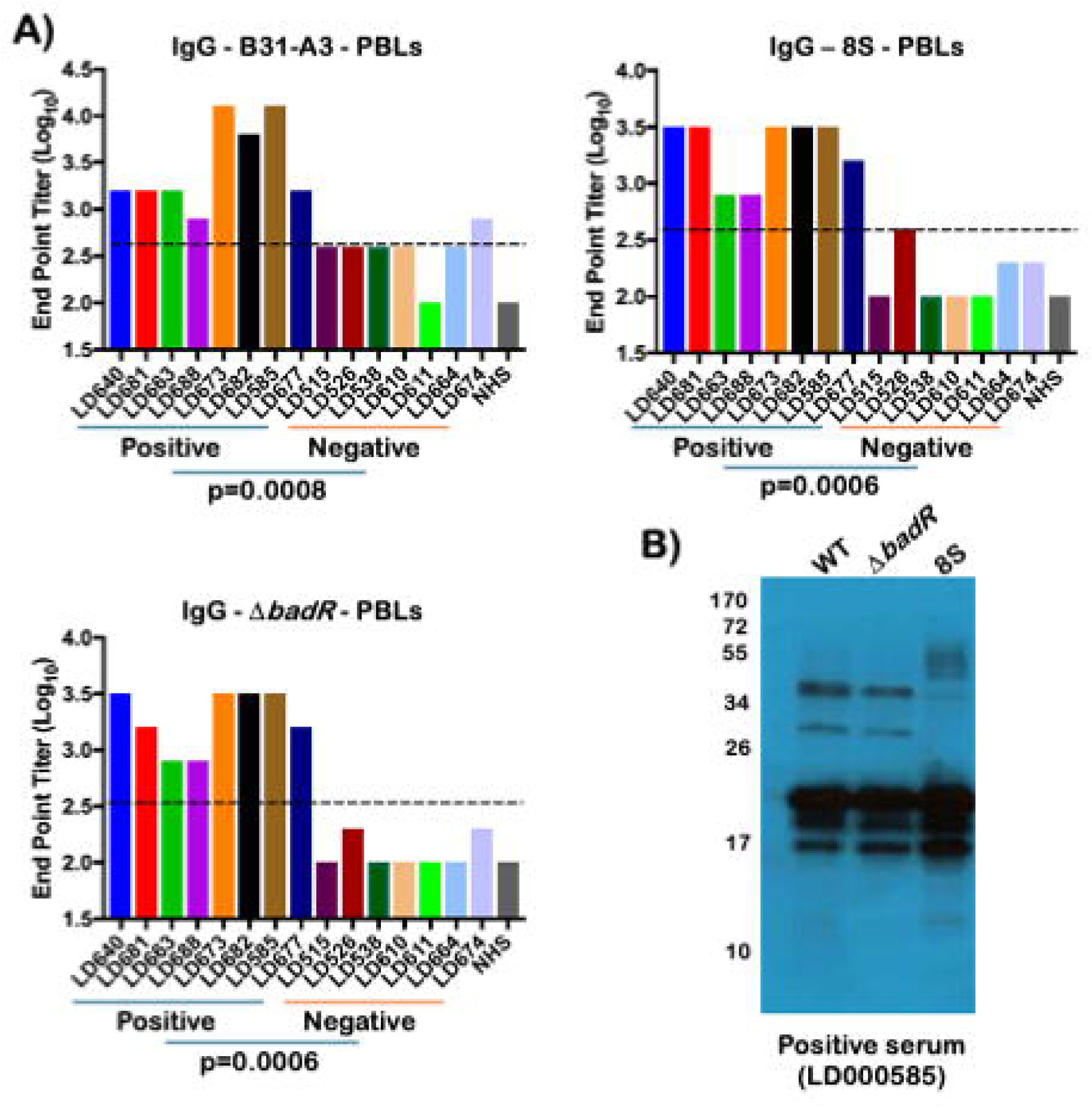
Reactivity of serum from Lyme disease patients with borrelial lipoproteins. 5A) Enzyme linked immunosorbent assay (ELISA) was performed with borrelial lipoproteins from B31-A3, Δ*badR* and 8S mutants coated as antigens on 96 well ELISA plates. Serial two-fold dilution of human serum samples (from 1:50 to 1:6400) from patients with or without Lyme disease (blind, no patient information provided by Lyme BioBank) were tested for reactivity. Plates were developed using anti-human IgG antibodies conjugated with horseradish peroxidase (HRPO) and o-phenylenediamine dihydrochloride (OPD) substrate solution and read at 450 nm using a microplate reader. Levels of IgG from 15 human serum samples are shown. Endpoint antibody titers of blinded human serum samples against lipoproteins from *Bb* strain B31-A3, 8S, and Δ*badR*. Normal human serum obtained from a local blood bank was used as a negative control. Cutoff lines indicate endpoint titers ≥2 serial dilution steps above the negative control, and samples exceeding this threshold were classified as high reactivity. Following unblinding, differences between positive and negative sample groups were analyzed using a Mann-Whitney U test; corresponding *p* values are shown at the bottom of each graph. A *p* value < 0.05 was considered statistically significant. 5B) Immunoblot analysis of a sample of human serum that exhibited high titers by ELISA. Proteins from lysates from *Bb* strain B31-A3, 8S and Δ*badR* were separated on a SDS-12.5% PAGE gel and transferred to PVDF membrane. Blots were developed using HRPO-conjugated anti human IgG and ECL system.

Among the isotypes of the human antibodies, IgG was predominant in several samples positive for Lyme disease, while the presence of IgM antibodies that reacted with PBLs were significant in positive serum samples (Fig. 6). Levels of IgG and IgG2 were significantly higher (p<0.05) against *Bb* lysate while IgM and IgG levels were significantly higher using PBLs as the coating antigens. The levels of significance were determined using a known *Bb*-negative human serum sample followed by comparison between positive and negative sample groups using a two-tailed Mann-Whitney U test. There were no significant differences in other isotypes between positive and negative human serum samples. Although the human serum samples were tested without prior clinical correlates, several serum samples that reacted with both *Bb* lysates or with PBLs correlated with clinical data compared retrospectively (Supplementary Table 1 and 2) (40). These results demonstrate that the Δ*badR* and 8S mutant strains retain immunologically relevant surface lipoproteins that are recognized by human antibodies, supporting their development as transmission-blocking biologics. Furthermore, this detailed isotype analysis provides insights into variations in human humoral responses against *Bb*, highlighting roles of IgM and IgG specific antibodies for developing efficacious formulations.

**Figure 6:**
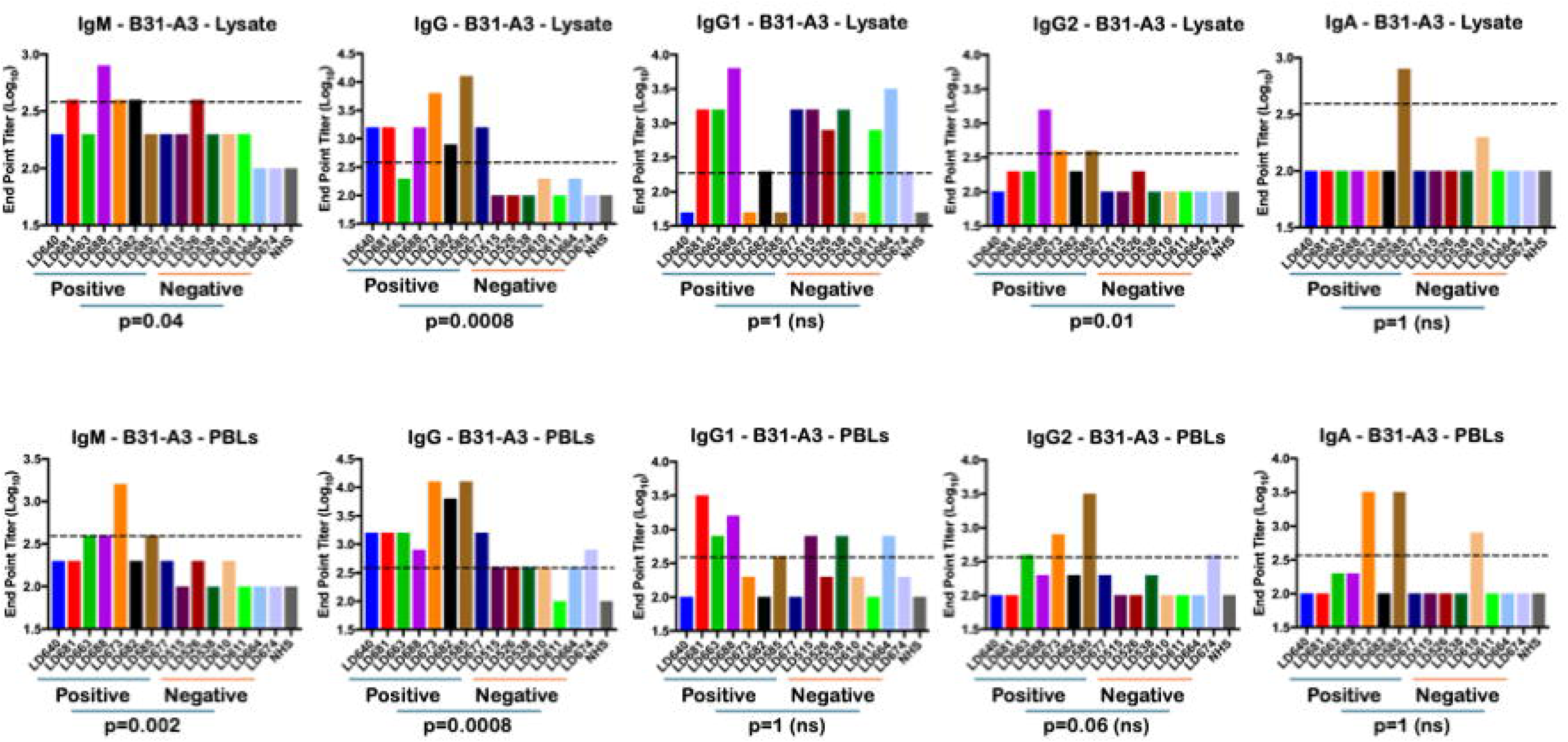
Isotype of human antibodies against *Bb* antigens. ELISA-based assays were performed using total lysates from *Bb* strain B31-A3, 8S and Δ*badR* to determine the isotype of the 15 human serum samples from patients with or without Lyme disease as described in Fig 5. Bar graphs showing end point titers of human serum samples determined using HRPO-conjugated secondary antibodies specific to human IgM, IgG, IgA, IgG1 and IgG2. Plates were read at 490 nm using a microplate reader. Levels of reactivity of normal human serum from the local blood bank served as a negative control. Cutoff lines indicate endpoint titers ≥2 serial dilution steps above the negative control, and samples exceeding this threshold were classified as high reactivity. Following unblinding, differences between positive and negative sample groups were analyzed using a Mann-Whitney U test; corresponding *p* values are shown at the bottom of each graph. A *p* value < 0.05 was considered statistically significant.

### Protective efficacy of purified lipoproteins from mutant borrelial strains

Since borrelial lipoproteins play a vital role both in the induction of protective immune responses as well as in the pathogenic mechanisms of *Bb*, supported by the presence of multiple outer surface proteins and significant antibody reactivity against human and mouse sera, we hypothesized that purified borrelial lipoproteins (PBLs) from the mutant strains are likely the major determinants influencing the protective capability of intact mutant spirochetes. Correlates of humoral immune response to intradermal inoculation of PBLs in C3H/HeN mice were established using an ELISA based method (Fig.7). The levels of IgG, IgG1, IgG2a and IgG2b were found to be significantly elevated compared to those in naïve controls at 28 dpv, while levels of IgM, IgA and IgG3 were not. To determine the ability of borrelial lipoproteins to confer protection against *Bb*, mice were immunized with PBLs from the parental *Bb* strain B31-A3 and mutant 8S and Δ*badR* strains. As shown in Table 2, C3H/HeN mice immunized via intradermal route with PBLs from B31-A3 and 8S were protected following challenge with *Bb* infected *I. scapularis* nymphs (confirmed via qPCR) with no viable spirochetes isolated from different infected tissues. The levels of protection conferred by PBLs from the Δ*badR* mutant were lower with two out of six mice positive for viable spirochetes in select tissues. Immunization with recombinant lipidated OspA, used as a positive control, was protective, with no tissues testing positive for *Bb*, whereas five of six unvaccinated control mice (PBS-treated) had tissues that were culture-positive for spirochetes.

**Figure 7:**
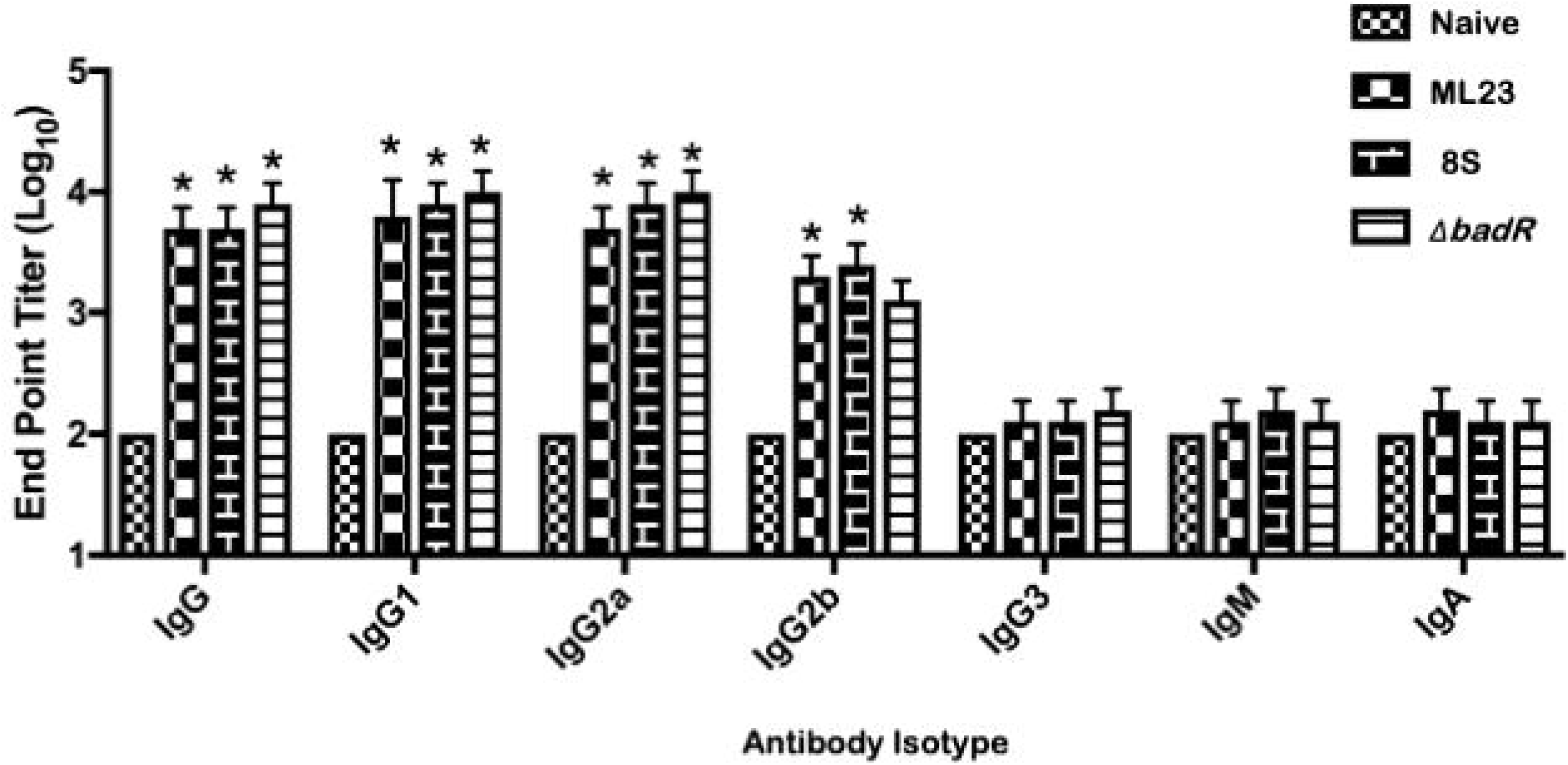
Isotype of antibodies induced in C3H/HeN mice at day 28 following vaccination with borrelial lipoproteins. Serum from mice vaccinated with lipoproteins from *Bb* strain B31-A3 ML23, 8S and Δ*badR* at day 0 and 14 were obtained 4 weeks after last vaccination and reactivity/isotype determined using total borrelial lysates from *Bb* strain B31-A3 using ELISA based method. Isotype specific mouse secondary antibodies conjugated to alkaline phosphatase (AP) and PNPP were used as substrate. Absorbance was measured at 405 nm with serum from naïve C3H/HeN mice serving as negative control. Note significant increase in levels of IgG, IgG1, IgG2a and IgG2b in vaccinated mice compared to naïve controls. The asterisks indicate significant difference between unvaccinated and vaccinated groups analyzed using a Mann-Whitney U test. A *p* value < 0.05 was considered statistically significant.

**Table 2:**
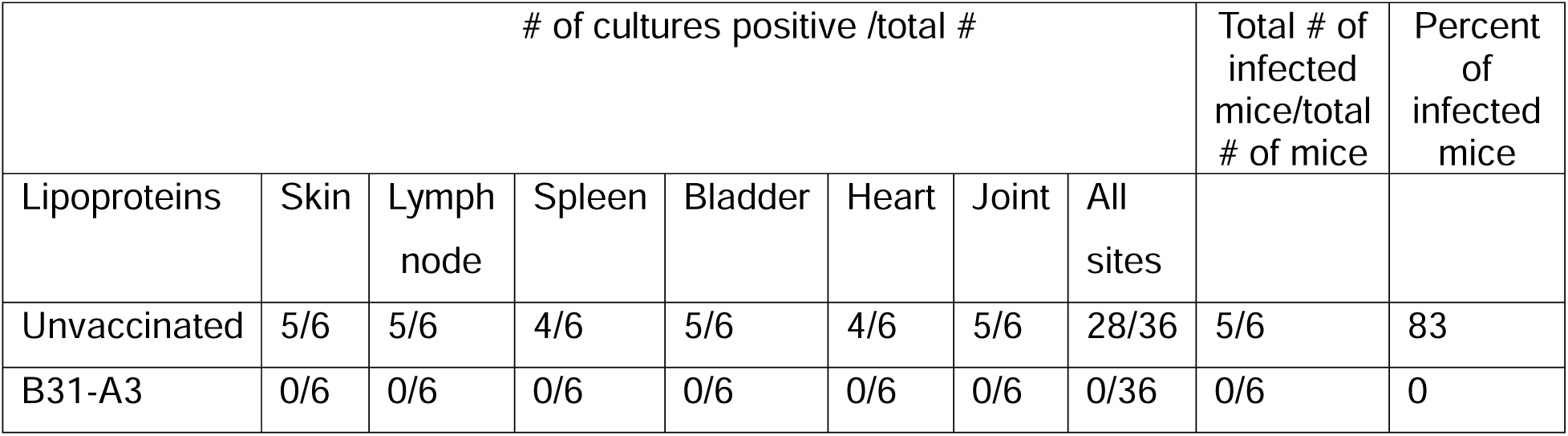
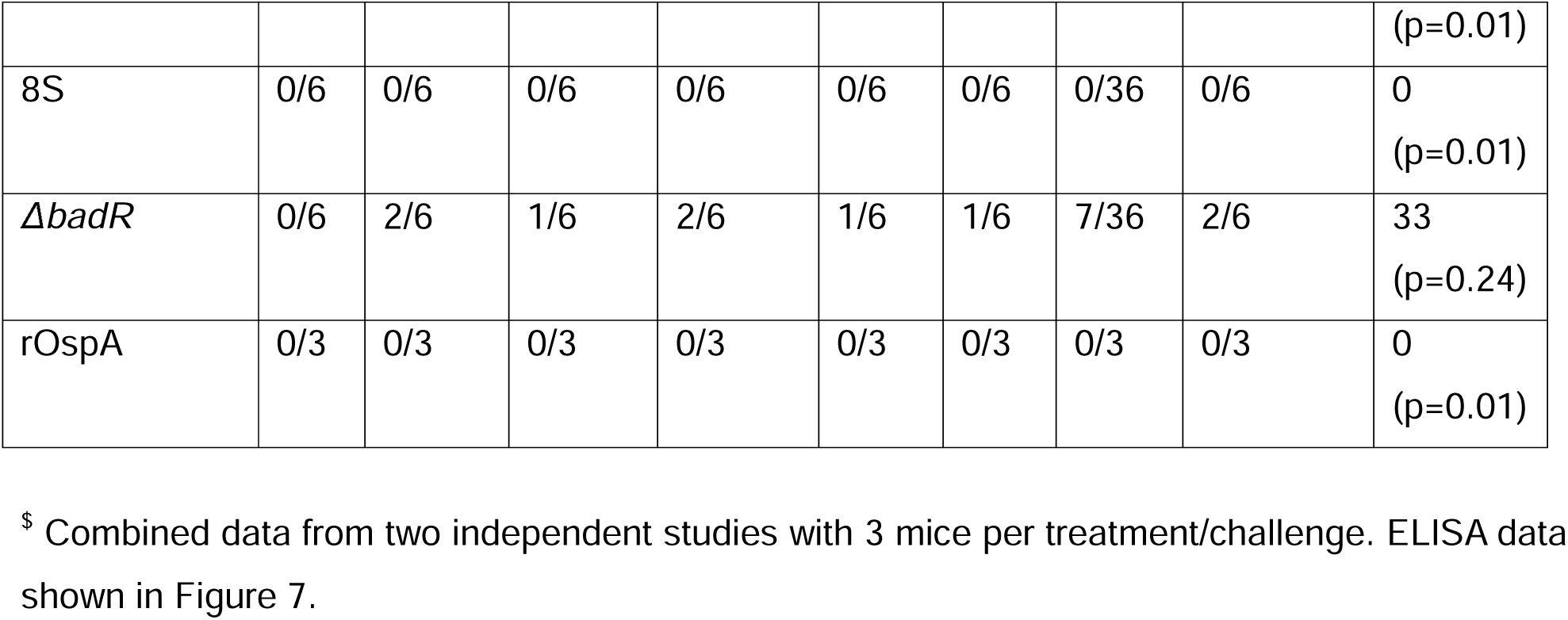
Efficacy of protection conferred by lipoproteins from mutant strains of *Bb* against tick challenge with *Bb* strain B31-A3^$^.

These data sets are representative of two independent studies with three mice each to facilitate using *Bb* infected nymphs obtained from a single infected C3H/HeN mice to ensure levels of *Bb* challenge are comparable between each treatment. Vaccination of C3H/HeN mice with PBLs from parental strain B31-A3 and 8S induced protective responses that resulted in a significant reduction (p=0.01) of *Bb* colonization following challenge with infected *Is* nymphs, serving as a key finding to advance strategies to develop reservoir host-targeted oral formulations.

## Discussion

There is a dire need to develop products and strategies to reduce the burden of tick-borne pathogens in reservoir hosts in the endemic areas where these diseases are prevalent to eliminate or reduce the incidence of tick-borne diseases in humans (30). Moreover, the increasing incidence and prevalence of Lyme disease across a wider geographic landscape underscore the urgent need for innovative approaches that disrupt the enzootic transmission cycle of the causative agent, *Borrelia burgdorferi* (30). Utilizing proteins from *Bb* expressed in heterologous bacterial hosts as antigens, several vaccines have been developed primarily targeting accidental hosts, such as humans (37, 41–43) and dogs (44–46). Several of these vaccination approaches focused on individual antigens such as Outer surface protein A or a combination of recombinant proteins (47–51). Although these strategies have shown varying efficacy, they rely on defined antigens are less likely to fully capture the antigenic complexity encountered during natural infection. Because the pathogen persists in nature through a complex cycle involving a variety of vertebrate reservoir hosts and *Ixodid* ticks, reservoir-targeted interventions have emerged as a promising strategy to reduce pathogen prevalence in ecosystems where the disease is endemic (7, 31). Current vaccine studies suggest that pathogen-derived antigenic repertoires that more closely mimic the physiological surface composition of the spirochete may provide broader immune recognition and improved transmission-blocking potential compared to recombinant antigens (52).

Recent expansion of a range of mutant strains of *Bb* either from isolation of clones lacking one or more plasmids or through targeted mutagenesis have opened avenues to explore the utility of these mutants for formulating biologics that limit pathogen survival in ticks or vertebrate hosts. A mutant lacking a surface exposed protein of *Bb* encoded by *bb405* that was expressed throughout the enzootic cycle had an essential *in vivo* function but with no detectable antibody responses in serum from natural infections but was capable of eliciting high and prolonged antibody titers protecting against tick-borne infection as a recombinant protein (51, 53). In addition, a *bba52 and bbi39* mutants lacking an ability to be transmitted between tick and mammalian hosts serve as a potential targets for blocking transmission of *Bb* underscoring the importance of targeted borrelial mutants as key products that can be leveraged to interfere with life cycle of Lyme spirochetes (50, 54). In this study, we investigated whether mutant strains of *Bb* that hyper-express surface lipoproteins could serve as sources of immunogenic lipoprotein complexes capable of disrupting the pathogen’s transmission cycle. The two mutants examined (8S and Δ*badR*) exhibit constitutive activation of the RpoS regulon, resulting in elevated expression of several mammalian-phase lipoproteins (24, 25, 55). The RpoS regulatory network plays a central role in enabling *Bb* to transition between tick and mammalian environments (56–58). In the Δ*badR* mutant, deletion of the transcriptional regulator BadR results in derepression of RpoS. Similarly, alanine substitutions of conserved residues of the Carbon Storage Regulator A of *Bb* (CsrA_Bb_) in the mutant strain (8S mutant) lead to increased expression of RpoS-regulated genes. Consequently, these mutants display enhanced expression of multiple outer surface lipoproteins normally induced during mammalian infection, but incapable of survival within the mammalian hosts, providing a unique opportunity to exploit these strains as live attenuated formulations as well as biological sources of antigenic determinants associated with host colonization and induction of host immune response to limit survival of *Bb* during different stages of enzootic cycle (59).

Vaccination studies in C3H/HeN mice demonstrated that intradermal immunization with live mutant strains elicited robust antibody responses and significantly reduced acquisition of *Bb* by naïve *Ixodes scapularis* larvae feeding on vaccinated hosts (Fig. 1 and 2), consistent with observations using other live attenuated vaccines (17). Because ticks acquire spirochetes during blood feeding, host antibodies can interact and limit spirochete survivability in conjunction with other host factors such as complement in the blood meal, thereby limiting pathogen acquisition by feeding larvae (60). In addition, OspA antibodies exhibit borreliacidal activities within the tick gut, reducing the level of spirochete transmission from infected nymphs to the mammalian hosts, thereby affecting the entire tick-mouse-tick cycle of spirochetes (61). Such antibody-mediated interference at the tick-host interface is a key mechanism underlying reservoir-targeted vaccination strategies (62). Our findings suggest that immunization with live attenuated strains produced variable levels of protection across studies, especially following challenge after an extended interval after two-dose immunization schedule (Fig. 2). Although live attenuated strains induced significantly higher antibody responses in vaccinated mice after intradermal inoculations and reduced pathogen acquisition by ticks, reduction in tissue colonization was less pronounced in some groups. There was a significant reduction in the levels of *Bb* acquired by naïve larvae from mice immunized with Δ*badR* compared to those immunized with ML23 or 8S, suggesting that levels of protection after a longer immunization schedule likely results in immune responses that are directed at additional antigens or that the nature of the antibody response likely limits *Bb* survival via complement dependent effects or increased memory T cell responses, among others. It is also possible that levels of hyperexpression of lipoproteins in the Δ*badR* mutant is less likely to mask additional immunoreactive proteins compared to the 8S strain that has a higher expression of OspB, a scenario that has been shown to occur in an engineered mutant *Bb* strain lacking OspA/B/C (19). In addition, this difference likely reflects a variety of other possibilities such as spirochetes from a higher challenge dose employed via needle challenge (10^5^ *Bb*/mouse) escaping less apparent differences in antibodies titers, withstand antibody mediated killing either due to antigenic variation or due to differences in antibody accessibility in different mammalian tissues (63). Spirochetes localized within connective tissue niches may be partially shielded from circulating antibodies, whereas organisms present in the bloodstream during tick feeding are more readily exposed to antibody-mediated mechanisms of neutralization (64, 65). These findings highlight the importance of distinguishing between sterilizing immunity and transmission-blocking immunity when evaluating vaccine strategies designed to disrupt the enzootic cycle. The protective immune response induced by the Δ*badR* mutant leading to a block in transmission of *Bb* to naïve ticks is a significant finding that will be further exploited to develop pathogen derived biologics to subvert the natural life cycle of *Bb*.

Consistent with these observations, culture-based analyses revealed a substantial reduction in recoverable spirochetes from tissues of mice vaccinated with the Δ*badR* mutant. The magnitude of protection also appeared to depend on the interval between times of vaccination and challenge, suggesting that peak antibody titers play an important role in limiting pathogen transmission to feeding ticks (66). These findings emphasize the importance of optimizing immunization schedules for reservoir-targeted interventions as an alternate to immunoprophylactic formulations for administration to accidental hosts such as humans or dogs. Since *B. burgdorferi* is primarily an extracellular pathogen, protective immunity largely depends on antibodies that recognize surface-exposed antigens and promote complement-mediated killing or opsonization (67). Vaccines based on purified surface antigens can therefore be highly effective in inducing neutralizing antibodies directed toward proteins either expressed at higher levels or are accessible during infection(68). Hence, the premise of targeting defined antigens such as OspA and OspC that are known to induce strong neutralizing antibody responses without exposing the host to live organisms is strong, there are caveats such as antigenic or strain specific variations of these individual lipoproteins influencing their protective efficacy (69, 70). Variations in host responses to these antigens are likely to play a role in the utility of these preparations. Moreover, recombinant versions of several borrelial lipoproteins such as OspA (32, 42, 43, 71–73), OspC (74–78), OspD (79, 80), OspE(81), BBK32(82, 83), LP6.6 (84) BBA64 (85, 86), BBA52(54), Lmp1 (87), and BBA34 (19, 20), among others, have been evaluated for their protective response either as single antigens or in combination (36, 80). We therefore hypothesized that purified lipoproteins, derived from the mutants that hyper-express a large array of these immunologically relevant lipoproteins (due to derepression of RpoS), are likely to provide enhanced protection and can be formulated for delivery via different routes of immunization.

To isolate these lipoproteins, spirochetal lysates were fractionated using Triton X-114, which enriched detergent-phase extracts for membrane-associated lipoproteins that exhibited strong immunoreactivity with infection-derived sera (39). The absence of the periplasmic flagellin FlaB confirmed the selectivity of this method (Fig. 3). Proteomic analysis identified several abundant lipoproteins including OspA, OspB, OspC, OspD, and LP6.6 (Fig. 4). These proteins are known to play important roles in both tick colonization and mammalian infection, indicating that the detergent-phase extracts capture a biologically relevant subset of surface antigens encountered during natural infection and host immune responses induced are likely to significantly decrease or block *Bb* acquisition by naïve larvae (84, 88, 89).

A notable observation was the increased abundance of OspB in the 8S mutant compared with the parental and Δ*badR* strains (Fig. 4). A number of prior studies have established a major role for OspB in antibody-dependent control of *Bb* with variation in the *ospB* gene, resulting in a generation of escape variants in response to an OspB monoclonal antibody (90–92). In addition, antibodies against OspB protect C3H/HeN mice from isolates that express truncated OspB antigens (93) and a multi-protein, chimeric immunogen that incorporates epitopes from OspB have been shown to elicit *Bb* killing via complement dependent and independent mechanisms (94). Immunoblot analyses also revealed a lower molecular-weight OspB-reactive species in the parental and Δ*badR* strains, suggesting proteolytic processing of this protein (Fig1B). In contrast, the 8S mutant predominantly expressed full-length OspB, indicating that regulatory pathways influenced by CsrA*_Bb_* may indirectly affect proteolytic processing of surface proteins. Although the functional significance of this observation remains unclear, proteolytic processing of outer surface proteins has been proposed as a mechanism by which *B. burgdorferi* modulates antigen exposure and immune recognition (93, 95). An advantage of using detergent-phase extracts from these mutants is that they preserve the relative abundance and stoichiometry of lipoproteins expressed in response to the host-specific environmental cues and to inactivate live spirochetes. Unlike cocktails of individually expressed recombinant proteins, these naturally derived antigenic complexes likely present multiple epitopes in configurations that more closely resemble those encountered during infection. This feature may promote broader antibody responses targeting multiple surface antigens simultaneously (39).

Analysis of immune recognition using mouse infection-derived sera and human serum samples demonstrated that these lipoprotein fractions contain multiple immunoreactive antigens that are recognized during natural infection with *Bb*. Several human serum samples exhibited strong IgM and IgG responses against both whole-cell lysates and lipoprotein extracts (Figures 5 and 6). These findings are consistent with the known kinetics of humoral responses observed during the course of Lyme disease, in which early IgM responses are followed by durable IgG responses that can persist for years after infection (40, 96). Recently, the frequencies and abundance of IgM, IgG and IgA specific to an immunodominant antigen of *Bb,* VlsE (Variable major protein like sequence, Expressed) was predictable between early and late Lyme arthritis stages of human Lyme disease with IgG4 isotype being a recognizable biomarker of Lyme arthritis patients (97). Since the present study included a variety of lipoproteins instead of a single major immunodominant antigen, we believe the IgM and IgG responses in general are reflective of the sequence of class switching noted in human serum samples. In human samples, variation in IgG subtypes suggests that natural infection induces different antibody responses across hosts, whereas immunization with a single antigen that elicits only one antibody subtype may be insufficient, highlighting the need for an approach that incorporates an array of antigens. In addition, the reactivity of positive human serum samples against PBLs was significantly higher that the reactivity with total *Bb* lysate providing an avenue for exploiting PBLs as diagnostic reagents for Lyme disease (Fig 6).

To determine whether purified lipoprotein complexes alone were sufficient to induce protective immunity, we evaluated detergent-phase lipoproteins derived from the parental and mutant strains. Notably, mice vaccinated with lipoproteins derived from the parental strain or the 8S mutant were significantly protected with no detectable spirochetes in various tissues examined following challenge with infected nymphs (Table 2). Intradermal immunization with these preparations induced strong IgG responses, predominantly IgG1, IgG2a, and IgG2b subclasses, indicating activation of mixed immune response involving both Th1- and Th2-associated immune pathways (98). It has been shown that reconstitution of B-cell deficient mice with normal mouse serum or polyclonal IgM but not IgG was able to reduce the spirochetal burden in ticks feeding on these mice suggesting natural IgM antibodies likely exert a borrelicidal effects on OspA-expressing spirochetes in tick-midgut (99). Moreover, it has been previously demonstrated that IgM responses that persist over a longer period of time had limited effects on bacterial dissemination or *Bb* burden contributing to reduced spirochete numbers in blood while the tissue penertrable IgG was bactericidal controlling Bb tissue burden (100). It is of importance to note that the levels of IgG, IgG1, IgG2a and IgG2b were elevated significantly in C3H/HeN mice vaccinated with PBLs serving as key immune correlate for utility of PBLs formulated as Bb transmission blocking biologics (Fig 7). These results demonstrate that naturally derived lipoprotein complexes can elicit robust protective immunity without the need for live spirochetes, especially through generating antibody subtypes that are key to limting Bb survival in tissues of mammalian hosts.

The protective efficacy observed here compares favorably with previously reported subunit vaccines targeting individual antigens such as OspA (69). While OspA-based vaccines primarily generate antibodies that kill spirochetes within the tick midgut during feeding (101), the lipoprotein complexes described in this study contain multiple antigens expressed during mammalian infection. As a result, they may induce broader immune responses capable of targeting spirochetes at multiple stages of the transmission cycle and reduce *Bb* survival within the vertebrate hosts (40).

These proof-of-concept findings are significant as they advance the development of pathogen-derived lipoprotein formulations as biologics for reservoir-targeted vaccination. Delivery through oral bait systems could provide a practical strategy for immunizing wild reservoir hosts in endemic regions. Similar approaches have been successfully implemented for other zoonotic pathogens, demonstrating the feasibility of large-scale environmental vaccination programs (30, 31, 47, 102, 103). Future studies applying such strategies could reduce pathogen persistence in natural cycles and ultimately decrease the chances of transmission of *Bb* via ticks, leading to an eventual decrease in the incidence of human Lyme disease. There are several caveats to the studies reported such as these experiments being conducted in a single laboratory mouse model and immunization regimens to establish immune parameters of protection following one or two doses. Therefore, additional studies are needed to evaluate efficacy in natural reservoir hosts such as *Peromyscus leucopus*. In addition, the durability and the mechanisms associated with the protective immune responses induced by the lipoproteins require further investigation (4).

In summary, mutant strains of *B. burgdorferi* with dysregulated surface antigen expression can serve as valuable sources of immunogenic lipoprotein complexes capable of interfering with pathogen transmission. By capturing the relative stoichiometry of diverse array of surface-exposed and immunogenic lipoproteins, these preparations generate broad antibody responses that limit pathogen acquisition by feeding ticks. This work establishes a framework for exploiting mutants as sources of biologically relevant antigenic repertoires for reservoir-targeted disease control strategies advancing similar studies utilizing a variety of mutants strains of *Bb* that are currently available for these purposes.

## Materials and Methods

### Animals and Ethics Statement

The animal facilities at The University of Texas at San Antonio (UTSA) are part of Laboratory Animal Resources Center (LARC), which is an AAALAC International Accredited Unit. 6-8-week- old female C3H/HeN and BALB/c mice (Charles River Laboratories, Wilmington, MA) were used in this study. All animal experiments were conducted following NIH guidelines for housing and care of laboratory animals and in accordance with protocols approved by the Institutional Animal Care and Use Committee of UTSA. Based on these guidelines, general condition and behavior of the animals were monitored by trained laboratory and LARC staff daily and methods to minimize pain and discomfort were adopted as needed in this study.

### Bacterial strains and growth conditions

A low passage infectious clonal isolate of *Borrelia burgdorferi* B31-A3 (104), mutant strains such as Δ*badR* (B31 isolate, *badR* (BB0693) deficient, Str^r^) (22), 8S (B31 isolate, CsrA*_Bb_* with 8 critical residues replaced with alanine, Gent^R^) (24) and ML23 (derived from strain *Bb*-B31 lacking linear plasmid 25 or lp25-deficeint strain from *Bb* B31-A3) (26, 27) were propagated at 32°C in liquid Barbour-Stoenner-Kelly (BSK-II, pH 7.6) media supplemented with 6% heat inactivated rabbit serum (Pel-Freez Biologicals, Rogers, AR) with appropriate antibiotics (Sigma-Aldrich, St. Louis, MO) as previously described (20, 22, 24, 25, 105, 106). Once the cultures reached a density between 1-2×10^7^ spirochetes /mL, viable spirochetes were enumerated by dark field microscopy and used for intradermal inoculation of C3H/HeN mice twice at a dose of 1×10^5^ *Bb* per mouse.

### Animal injections and tick larvae feeding

For infection studies, *Bb* cultures were verified by PCR for the presence of the virulence plasmids lp25 and lp28-1 on the day of injection. Following confirmation, cultures were centrifuged at 4,000 × g for 20 min at 4 °C, and the pellets were washed three times with sterile HBSS (4,000 × g for 5 min at 4 °C). The cells were then resuspended in BSK-II medium supplemented with 6% heat-inactivated rabbit serum to a final concentration of 1 × 10J spirochetes/mL. Each mouse received an intradermal injection of 100 µL of the spirochete suspension (83, 107–110).

Twenty-eight days after needle challenge, the dorsal skin of the mice was shaved, and sterile plastic feeding capsules were affixed to the skin using a non-abrasive adhesive and was allowed to dry overnight. *Ixodes scapularis* larvae were obtained from the tick rearing facility at Oklahoma State University (Stillwater, OK) and maintained at 22 °C with 90% humidity under a 15-h light/9-h dark cycle. 50-100 naïve larvae were placed inside each capsule, which was then sealed with perforated caps to facilitate tick attachment and feeding. After 3-7 days, engorged larvae were collected and *Bb* burden in ticks were assessed. Briefly, four engorged larvae were crushed and analyzed for spirochete burden. Total genomic DNA was extracted and analyzed by quantitative real-time PCR using primers specifically *Bb flaB*. Spirochete burden was normalized to tick actin and reported as relative copy numbers of *Bb_flaB* to *I. scapularis_actin*. The remaining larvae were placed in sterile tubes covered with perforated parafilm and incubated at 23 °C under humid conditions for 5-8 weeks to allow for molting into nymphs. All mouse infection studies were conducted in a Biosafety Level 2 facility.

### Tick challenge

To generate *Bb* infected nymphs for challenge studies, naïve larvae were allowed to feed on C3H/HeN mice that had been infected intradermally with 1 × 10J *B. burgdorferi* strain B31-A3 as previously described (107). To assess spirochete acquisition in flat nymphs, five nymphs from the stock tubes were crushed individually and analyzed for spirochete burden, as mentioned in the previous paragraph and reported as relative copy numbers of *Bb_flaB* to *I. scapularis_actin* (Supplementary Fig. 1). After confirming the *Bb* load, infected nymphs were then used to challenge naïve or immunized C3H/HeN mice (immunized with purified lipoproteins from different mutant strains or recombinant OspA as a control) by allowing ticks to feed to repletion (3-5 days).

### Necropsy and quantification of *Bb* burden in tissues

Twenty-eight days after needle challenge or tick challenge, mice were euthanized by carbon dioxide inhalation followed by cardiac puncture, serving as terminal bleed and as secondary method of euthanasia. Skin, lymph nodes, spleen, heart, bladder and joint tissues were aseptically isolated and inoculated into BSK-II medium, blindly passaged at day 5, and scored for growth by examining cultures using dark field microscopy at day 14 post-isolation in primary and blind-pass cultures (107, 109, 110). Another set of skin, lymph nodes, spleen and joint samples were stored at −80°C for genomic DNA extraction and qPCR analysis.

### Genomic DNA extraction and qPCR

Genomic DNA from the skin, spleen and lymph nodes from mice as well as from larvae and nymphs were extracted using High Pure PCR Template Preparation Kit (Roche, Indianapolis, IN). Samples were lysed with 200 µL of lysis buffer in a BeadBlaster™ 24 microtube homogenizer using Zirconia beads. Further, 40 µL of proteinase K (20 mg/mL) and 20 µL of Collagenase (1 mg/mL) was added and incubated overnight at 57°C. Next, DNA was extracted as directed by the manufacturer using the column. *Bb* burden in the samples was determined by a SYBR-based qPCR assay (PowerUp SYBR Master Mix, Applied Biosystems) using *Bb*FlaB gene specific primers and normalized to tick actin or mouse actin and the reaction was performed in a StepOne Plus real time PCR system (Applied Biosystems). Standard curves for all three genes were performed by full length genes cloned into pCR2.1 plasmid and the results are presented as the ratio of *flaB* to mouse or tick actin copy number.

### Cloning and overexpression of OspA

Genomic DNA from *Bb* strain B31-A3 was prepared and *ospA* was amplified using primers OspA-F/Xba1- ACGC TCTAGA ATGAAAAAATATTTATTGGGAATAGGTCTA and OspA-R/Sal1 ACGCGTCGACTTATTTTAAAGCGTAATTAATTTCATCAAG. The amplicon was first cloned into pCR2.1 and transformed into *E. coli* TOP10 cells, exploiting the blue/white screening method. The insert was excised using the engineered restriction enzyme sites and cloned into expression plasmid pMAL-2x. Plasmids with the right insert were identified using pMAL2c specific primers flanking the insert and forward and were transformed into *E. coli* RosettaTM (Novagen). OspA fused to Maltose binding protein was induced with IPTG and purified using amylose beads as described previously (107, 110–112). Purified OspA was quantified using BCA and stored at −20°C in aliquots until used. A similar construct was generated using pET23a vector and OspA fused to a 6X-HisTag was purified using NiNTA beads as described previously (20).

### Lipoprotein purification from *Bb* strains

Lipoproteins were extracted from the wildtype and mutant strains as previously described (85, 113). Briefly, 1×10^9^ *Bb* was solubilized in 1 mL of PBS (Phosphate Buffered Saline, pH7.4) containing 1% Triton^®^ X-114 (TX-114) (Sigma Aldrich) by gentle rocking at 4°C overnight. The pellet containing the TX-114 insoluble material was removed by two centrifugations at 15,000xg at 4°C for 15 mins. The supernatant was transferred to a sterile tube and incubated at 37°C for 15 mins. Then the mixture was centrifuged at 15,000xg for 15 mins at RT. The top aqueous phase was transferred to a new tube and re-extracted one more time with 1% TX-114 as described above. The bottom detergent phase was washed thrice with 1 mL PBS. The final detergent phase proteins were precipitated by adding 10-fold volume of ice-cold acetone, precipitates were collected by centrifugation at 15,000xg at 4°C for 30 mins, acetone was removed by drying, and proteins were resuspended in PBS. Insoluble (Pellet), soluble (aqueous phase) and lipoprotein (TX-114 phase) fractions were analyzed by SDS-PAGE gel. Purified Borrelial Lipoproteins (PBLs) were quantified using a BCA assay kit (Thermo Scientific) and stored at −20°C until further use.

### Proteomic analysis of PBLs

PBLs isolated from infectious B31 strain (WT), 8S and Δ*badR* mutant strains were submitted in triplicate to UTMB Proteomics facility for Mass Spec analysis to determine the composition of the detergent phase proteins. Each sample mixture was solubilized with 5% SDS, 50 mM triethyl ammonium bicarbonate, pH 7.55 in a final volume of 25 μL. The sample was then centrifuged at 17,000 g for 10 min to remove any debris. Proteins were reduced using a solution of 20 mM Tris-2-carboxyethyl phosphine (TCEP, Thermo, Scientific catalog #77,720) and incubated at 65 °C for 30 min. The sample was cooled to room temperature and 1 μL of 0.5 M iodoacetamide acid added and allowed to react for 20 mins in the dark. 2.75 μl of 12% phosphoric acid was added to the protein solution. 165 μL of binding buffer (90% Methanol, 100 mM TEAB final; pH 7.1) was then added to the solution. The resulting solution was added to S-Trap spin column (protifi.com) and passed through the column using a bench top centrifuge (30 s spin at 4000×g). The spin column was washed with 400 μL of binding buffer and centrifuged. The binding buffer wash was then repeated two more times. Trypsin was added to the protein mixture in a ratio of 1:25 in 50 mM TEAB, pH 8, and incubated at 37 °C for 4 h. Peptides were eluted with 75 μl of 50% acetonitrile, 0.2% formic acid, and then washed again with 75 μl of 80% acetonitrile, 0.2% formic acid. The combined peptide solution was then dried in a speed vac and resuspended in 2% acetonitrile, 0.1% formic acid, 97.9% water and placed in an autosampler vial.

### NanoLC MS/MS Analysis

Peptide mixtures were analyzed by nanoflow liquid chromatography-tandem mass spectrometry (nanoLC-MS/MS) using a nano-LC chromatography system (UltiMate 3000 RSLCnano, Dionex), coupled on-line to a Thermo Orbitrap Fusion mass spectrometer (Thermo Fisher Scientific, San Jose, CA) through a nanospray ion source (Thermo Scientific). A trap and elute method was used. The trap column was a C18 PepMap100 (300 um X 5 mm, 5 um particle size, Thermo Scientific) and the analytical column was an Acclaim PepMap 100 (75 um X 25 cm, Thermo Scientific). After equilibrating the column in 98% solvent A (0.1% formic acid in water) and 2% solvent B (0.1% formic acid in acetonitrile (ACN)), the samples (1 µL in solvent A) were injected onto the trap column and subsequently eluted (300 nL/min) by gradient elution onto the C18 column as follows: isocratic at 2% B, 0-5 min; 5-6 min 2%-4% B, 4% to 32% B, 5- 120 min; 32% to 90% B, 120-123 min; isocratic at 90% B, 123-126 min; 90% to 2%, 126-129 min; isocratic at 2% B, 129-130 min; 2% to 90% B, 130-134 min; isocratic at 90% B, 134-137 min; 90% to 2%, 137-140 min; and isocratic at 2% B, 140-145 min.

All LC-MS/MS data were acquired using Xcalibur, version 4.4.16.14 (Thermo Fisher Scientific) in positive ion mode using a top speed data-dependent acquisition (DDA) method with a 3 sec cycle time. The survey scans (m/z 375-1500) were acquired in the Orbitrap at 120,000 resolution (at m/z = 400) in profile mode, with maximum injection mode set to Auto and a normalized AGC target of 100%. The S-lens RF level was set to 60. Isolation was performed in the quadrupole with a 1.6 Da isolation window, and CID MS/MS acquisition was performed in centroid mode with detection in the ion trap (Rapid scan rate) with the following settings: CID collision energy = 30%; maximum injection time mode set to Dynamic; normalized AGC target of 20%. Monoisotopic precursor selection (MIPS) and charge state filtering were on, with charge states 2-7 included. Dynamic exclusion was used to remove selected precursor ions, with a +/-10 ppm mass tolerance, for 30 sec after acquisition of one MS/MS spectrum.

### Database Searching

Tandem mass spectra were extracted, and charge state deconvolved by Proteome Discoverer (Thermo Fisher, version 2.5.0.402). All MS/MS spectra were searched against a concatenated FASTA database which included a Uniprot *Bb* database using Sequest. Searches were performed with a parent ion tolerance of 10 ppm, and a fragment ion tolerance of 0.60 Da. Trypsin was specified as the enzyme, allowing for two missed cleavages. A fixed modification of carbamidomethyl (C) and variable modifications of oxidation (M) and deamidation of asparagine and glutamine, were specified in Sequest. Percolator was used to filter the peptide identification results to a false discovery rate of 1%. Label-free quantitation was performed using Minora. Differential expression analysis of sample groups was performed using the Precursor Ions Quantifier node in Proteome Discoverer, with protein abundance ratios calculated using a Pairwise Ratio scheme while hypothesis testing was performed using a t-test of the background population of proteins and peptides. The pathways associated with the identified proteins were determined by KEGG pathway search tool (https://www.genome.jp/kegg/mapper/search.html), against *Borrelia burgdorferi* B31 datasets.

### Immunization with lipoproteins or live bacterial strains

For immunization, mutant strains grown in BSK-II media were centrifuged at 4000x g for 20 minutes at 4°C and the pellets were washed three times with sterile HBSS at 4000x g for 5 minutes at 4°C and the cells were resuspended in BSK-II medium supplemented with 6% heat inactivated rabbit serum at a final concentration of 10^6^ spirochetes/mL as described previously (83, 108, 109, 111). Each mouse was injected with 100 µL of spirochetes via intra-dermal injections on day 0 and day 16 or 18. Similarly, purified lipoproteins or recombinant OspA were injected at a concentration of 100 µg/100 µL via intra-dermal injections on day 0 and day 14.

### Enzyme Linked Immunosorbent Assay (ELISA)

To determine the peripheral antibody levels, blood was collected from the lateral saphenous vein using Microvette CB 300 capillary tubes (Sarstedt AG & Co.KG), centrifuged at 4000xg for 10 minutes and the supernatant were collected and stored at −20°C until further use. ELISA was performed against borrelial lysate or PBLs as required. All the assays were performed using a Costar High binding Assay Plate (Corning). 10 µg of the antigen were resuspended in 10 mL of coating carbonate buffer (0.05M, pH 9.6) and 100 µL of coating antigen were added to each well and incubated overnight at 4°C. After washing three times with Tris Buffered Saline supplemented with Tween 20 (TBST), 100 µL of 1% Bovine Serum Albumin (BSA) (Probumin, Millipore) in TBST was added to each well and incubated for 1 hr at room temperature and washed three times with TBST. Serum samples were diluted in TBST starting from 1:50 up to 1:6400 dilutions and 100 µL of each diluted sample was added. After two hours of incubation at room temperature, the plates were washed five times with TBST and 100 µL of 1:3000 diluted secondary anti-mouse and anti-human IgG antibodies conjugated with Horseradish peroxidase (HRP) were added and incubated for 1 hr. After four washes, 100 µL of OPD substrate solution was added and read at 450 nm using Spark 10M (Tecan) microplate reader. In addition, plates were also developed using anti-mouse isotype specific antibodies conjugated to alkaline phosphatase. Levels of significance were determined using unpaired Student *t*-test using GraphPad Prism 7.0 and indicated in the respective figure legends.

### Immunoblot Analysis

Immunoblot analysis was performed to determine the antigenic profile of lipoproteins against mouse and human serum (blinded samples provided by Lyme disease Bio Bank). Equal amounts of purified lipoproteins (5 µg/sample) were separated using 12% polyacrylamide gels and then transferred to a polyvinylidene difluoride (PVDF) membrane, blocked with blocking buffer (10% non-fat dry milk in Tris buffer (pH 7.5) containing 200 mM Tris, 1.38 M NaCl, and 0.1% Tween 20) overnight and then incubated with infected serum collected from mouse or human patients or rFlaB immunized mouse serum at room temperature for 1 hr in blocking buffer, and treated with horseradish peroxidase (HRP) conjugated Goat anti-mouse or Goat anti-human IgG secondary antibody for 1 hr at room temperature. Immunoblots were developed using an enhanced chemiluminescence system followed by exposure to X-ray film.

### Statistical analysis

To test the vaccine-induced protection, animals were classified as culture-positive if at least one tissue yielded a positive culture result and culture-negative if all tested tissues were negative. The variance in protective efficacy was compared between vaccinated and unvaccinated groups using a two-sided Fisher’s exact test (17). This test is appropriate for small sample sizes and binary outcomes. A *p* value < 0.05 was considered statistically significant.

For human serum ELISA, endpoint titers were determined for all samples. Because only a single known negative control serum was available, blinded samples were interpreted relative to this control. Samples exhibiting endpoint titers ≥2 serial dilution steps above the normal human serum (indicated by cutoff lines in the figures) were classified as high reactivity. Comparisons between positive and negative sample groups were performed using a two-tailed Mann-Whitney U test. For mouse serum ELISA, vaccinated samples were compared with the unvaccinated samples using a two-tailed Mann-Whitney U test. A *p* value < 0.05 was considered statistically significant.

For tissue and larval genomic DNA analyses, differences between vaccinated and unvaccinated groups were evaluated using a two-tailed Student’s *t*-test. A *p* value < 0.05 was considered statistically significant. All analyses were performed in GraphPad Prism.

## Acknowledgements

This study was partly supported by Public Health Service grant 5R01AI152233 from the National Institute of Allergy and Infectious Diseases (JS), Bay Area Lyme Foundation (JS), pre-doctoral fellowships from the South Texas Center for Emerging Infectious Diseases (TMI), and the Center of Excellence in Infection Genomics (TCS). TCS was supported by a pre-doctoral fellowship from The Brown Foundation. JFSL is supported by the UTSA Graduate School and The National Institute of Allergy and Infectious Diseases of the National Institutes of Health under Award Number T32AI184340. The contents are solely the responsibility of the authors and do not necessarily represent the official views of the National Institutes of Health.

## References

1. Kugeler KJ, Earley A, Mead PS, Hinckley AF. 2024. Surveillance for Lyme Disease After Implementation of a Revised Case Definition - United States, 2022. MMWR Morb Mortal Wkly Rep 73:118–123.

2. Mead P, Hinckley A, Kugeler K. 2024. Lyme Disease Surveillance and Epidemiology in the United States: A Historical Perspective. J Infect Dis 230:S11–S17.

3. Manley W, Tran T, Prusinski M, Brisson D. 2025. Comparative ecological analysis and predictive modeling of tick-borne pathogens. J Med Entomol 62:199–206.

4. Bobe JR, Jutras BL, Horn EJ, Embers ME, Bailey A, Moritz RL, Zhang Y, Soloski MJ, Ostfeld RS, Marconi RT, Aucott J, Ma’ayan A, Keesing F, Lewis K, Ben Mamoun C, Rebman AW, McClune ME, Breitschwerdt EB, Reddy PJ, Maggi R, Yang F, Nemser B, Ozcan A, Garner O, Di Carlo D, Ballard Z, Joung HA, Garcia-Romeu A, Griffiths RR, Baumgarth N, Fallon BA. 2021. Recent Progress in Lyme Disease and Remaining Challenges. Front Med (Lausanne) 8:666554.

5. Radolf JD, Caimano MJ, Stevenson B, Hu LT. 2012. Of ticks, mice and men: understanding the dual-host lifestyle of Lyme disease spirochaetes. Nat Rev Microbiol 10:87–99.

6. Scheckelhoff MR, Telford SR, Hu LT. 2006. Protective efficacy of an oral vaccine to reduce carriage of Borrelia burgdorferi (strain N40) in mouse and tick reservoirs. Vaccine 24:1949–57.

7. Vannier E, Richer LM, Dinh DM, Brisson D, Ostfeld RS, Gomes-Solecki M. 2023. Deployment of a Reservoir-Targeted Vaccine Against Borrelia burgdorferi Reduces the Prevalence of Babesia microti Coinfection in Ixodes scapularis Ticks. J Infect Dis 227:1127–1131.

8. Bourgeois JS, Hu LT. 2024. Hitchhiker’s Guide to Borrelia burgdorferi. J Bacteriol 206:e0011624.

9. Strnad M, Rudenko N, Rego ROM. 2023. Pathogenicity and virulence of Borrelia burgdorferi. Virulence 14:2265015.

10. Stevenson B. 2023. The Lyme disease spirochete, Borrelia burgdorferi, as a model vector-borne pathogen: insights on regulation of gene and protein expression. Curr Opin Microbiol 74:102332.

11. Sadziene A, Thomas DD, Bundoc VG, Holt SC, Barbour AG. 1991. A flagella-less mutant of Borrelia burgdorferi. Structural, molecular, and in vitro functional characterization. J Clin Invest 88:82–92.

12. Sadziene A, Thompson PA, Barbour AG. 1996. A flagella-less mutant of Borrelia burgdorferi as a live attenuated vaccine in the murine model of Lyme disease. J Infect Dis 173:1184–93.

13. Kurtti TJ, Munderloh UG, Hughes CA, Engstrom SM, Johnson RC. 1996. Resistance to tick-borne spirochete challenge induced by Borrelia burgdorferi strains that differ in expression of outer surface proteins. Infect Immun 64:4148–53.

14. Sadziene A, Barbour AG, Rosa PA, Thomas DD. 1993. An OspB mutant of Borrelia burgdorferi has reduced invasiveness in vitro and reduced infectivity in vivo. Infect Immun 61:3590–6.

15. Sadziene A, Thomas DD, Barbour AG. 1995. Borrelia burgdorferi mutant lacking Osp: biological and immunological characterization. Infect Immun 63:1573–80.

16. Stafford KC, 3rd, Williams SC, van Oosterwijk JG, Linske MA, Zatechka S, Richer LM, Molaei G, Przybyszewski C, Wikel SK. 2020. Field evaluation of a novel oral reservoir-targeted vaccine against Borrelia burgdorferi utilizing an inactivated whole-cell bacterial antigen expression vehicle. Exp Appl Acarol 80:257–268.

17. Hahn BL, Padmore LJ, Ristow LC, Curtis MW, Coburn J. 2016. Live Attenuated Borrelia burgdorferi Targeted Mutants in an Infectious Strain Background Protect Mice from Challenge Infection. Clin Vaccine Immunol 23:725–31.

18. Curtis MW, Hahn BL, Zhang K, Li C, Robinson RT, Coburn J. 2018. Characterization of Stress and Innate Immunity Resistance of Wild-Type and Deltap66 Borrelia burgdorferi. Infect Immun 86.

19. Yas OB, Coleman AS, Lipman RM, Sharma K, Raghunandanan S, Alanazi F, Rana VS, Kitsou C, Yang X, Pal U. 2024. A systemic approach to identify non-abundant immunogenic proteins in Lyme disease pathogens. mSystems 9:e0108723.

20. Raju BV, Esteve-Gassent MD, Karna SL, Miller CL, Van Laar TA, Seshu J. 2011. Oligopeptide permease A5 modulates vertebrate host-specific adaptation of Borrelia burgdorferi. Infect Immun 79:3407–20.

21. Kariu T, Sharma K, Singh P, Smith AA, Backstedt B, Buyuktanir O, Pal U. 2015. BB0323 and novel virulence determinant BB0238: Borrelia burgdorferi proteins that interact with and stabilize each other and are critical for infectivity. J Infect Dis 211:462–71.

22. Miller CL, Karna SL, Seshu J. 2013. Borrelia host adaptation Regulator (BadR) regulates rpoS to modulate host adaptation and virulence factors in Borrelia burgdorferi. Mol Microbiol 88:105–24.

23. Ouyang Z, Zhou J. 2015. BadR (BB0693) controls growth phase-dependent induction of rpoS and bosR in Borrelia burgdorferi via recognizing TAAAATAT motifs. Mol Microbiol 98:1147–67.

24. Karna SL, Prabhu RG, Lin YH, Miller CL, Seshu J. 2013. Contributions of environmental signals and conserved residues to the functions of carbon storage regulator A of Borrelia burgdorferi. Infect Immun 81:2972–85.

25. Karna SL, Sanjuan E, Esteve-Gassent MD, Miller CL, Maruskova M, Seshu J. 2011. CsrA modulates levels of lipoproteins and key regulators of gene expression critical for pathogenic mechanisms of Borrelia burgdorferi. Infect Immun 79:732–44.

26. Labandeira-Rey M, Skare JT. 2001. Decreased infectivity in Borrelia burgdorferi strain B31 is associated with loss of linear plasmid 25 or 28-1. Infect Immun 69:446–55.

27. Labandeira-Rey M, Seshu J, Skare JT. 2003. The absence of linear plasmid 25 or 28-1 of Borrelia burgdorferi dramatically alters the kinetics of experimental infection via distinct mechanisms. Infect Immun 71:4608–13.

28. Purser JE, Lawrenz MB, Caimano MJ, Howell JK, Radolf JD, Norris SJ. 2003. A plasmid-encoded nicotinamidase (PncA) is essential for infectivity of Borrelia burgdorferi in a mammalian host. Mol Microbiol 48:753–64.

29. Kitsou C, Pal U. 2022. Vaccines Against Vector-Borne Diseases. Methods Mol Biol 2411:269–286.

30. Gomes-Solecki M, Arnaboldi PM, Backenson PB, Benach JL, Cooper CL, Dattwyler RJ, Diuk-Wasser M, Fikrig E, Hovius JW, Laegreid W, Lundberg U, Marconi RT, Marques AR, Molloy P, Narasimhan S, Pal U, Pedra JHF, Plotkin S, Rock DL, Rosa P, Telford SR, Tsao J, Yang XF, Schutzer SE. 2020. Protective Immunity and New Vaccines for Lyme Disease. Clin Infect Dis 70:1768–1773.

31. Gomes-Solecki MJ, Brisson DR, Dattwyler RJ. 2006. Oral vaccine that breaks the transmission cycle of the Lyme disease spirochete can be delivered via bait. Vaccine 24:4440–9.

32. Steere AC, Sikand VK, Meurice F, Parenti DL, Fikrig E, Schoen RT, Nowakowski J, Schmid CH, Laukamp S, Buscarino C, Krause DS. 1998. Vaccination against Lyme disease with recombinant Borrelia burgdorferi outer-surface lipoprotein A with adjuvant. Lyme Disease Vaccine Study Group. N Engl J Med 339:209–15.

33. Yang XF, Pal U, Alani SM, Fikrig E, Norgard MV. 2004. Essential role for OspA/B in the life cycle of the Lyme disease spirochete. J Exp Med 199:641–8.

34. Dowdell AS, Murphy MD, Azodi C, Swanson SK, Florens L, Chen S, Zückert WR. 2017. Comprehensive Spatial Analysis of the Borrelia burgdorferi Lipoproteome Reveals a Compartmentalization Bias toward the Bacterial Surface. J Bacteriol 199.

35. Haake DA, Zuckert WR. 2018. Spirochetal Lipoproteins in Pathogenesis and Immunity. Curr Top Microbiol Immunol 415:239–271.

36. Brown EL, Kim JH, Reisenbichler ES, Höök M. 2005. Multicomponent Lyme vaccine: three is not a crowd. Vaccine 23:3687–96.

37. Dattwyler RJ, Gomes-Solecki M. 2022. The year that shaped the outcome of the OspA vaccine for human Lyme disease. NPJ Vaccines 7:10.

38. Chen S, Kumru OS, Zuckert WR. 2011. Determination of Borrelia surface lipoprotein anchor topology by surface proteolysis. J Bacteriol 193:6379–83.

39. Rao TD, Frey AB. 1998. Soluble Proteins Isolated fromBorrelia burgdorferiby Extraction with Triton X-114 Confer Resistance to Experimental Infection. Clinical Immunology and Immunopathology 89:94–104.

40. Ghosh R, Joung HA, Goncharov A, Palanisamy B, Ngo K, Pejcinovic K, Krockenberger N, Horn EJ, Garner OB, Ghazal E, O’Kula A, Arnaboldi PM, Dattwyler RJ, Ozcan A, Di Carlo D. 2024. Rapid single-tier serodiagnosis of Lyme disease. Nat Commun 15:7124.

41. de Silva AM, Zeidner NS, Zhang Y, Dolan MC, Piesman J, Fikrig E. 1999. Influence of outer surface protein A antibody on Borrelia burgdorferi within feeding ticks. Infect Immun 67:30–5.

42. de Silva AM, Fish D, Burkot TR, Zhang Y, Fikrig E. 1997. OspA antibodies inhibit the acquisition of Borrelia burgdorferi by Ixodes ticks. Infect Immun 65:3146–50.

43. de Silva AM, Telford SR, 3rd, Brunet LR, Barthold SW, Fikrig E. 1996. Borrelia burgdorferi OspA is an arthropod-specific transmission-blocking Lyme disease vaccine. J Exp Med 183:271–5.

44. Marconi RT, Garcia-Tapia D, Hoevers J, Honsberger N, King VL, Ritter D, Schwahn DJ, Swearingin L, Weber A, Winkler MTC, Millership J. 2020. VANGUARD(R)crLyme: A next generation Lyme disease vaccine that prevents B. burgdorferi infection in dogs. Vaccine X 6:100079.

45. Izac JR, O’Bier NS, Oliver LD, Jr., Camire AC, Earnhart CG, LeBlanc Rhodes DV, Young BF, Parnham SR, Davies C, Marconi RT. 2020. Development and optimization of OspC chimeritope vaccinogens for Lyme disease. Vaccine 38:1915–1924.

46. Chambers GZ, Chambers KMF, Marconi RT. 2024. A single immunization of Borreliella burgdorferi-infected mice with Vanguard crLyme elicits robust antibody responses to diverse strains and variants of outer surface protein C. Infect Immun 92:e0039624.

47. Bhattacharya D, Bensaci M, Luker KE, Luker G, Wisdom S, Telford SR, Hu LT. 2011. Development of a baited oral vaccine for use in reservoir-targeted strategies against Lyme disease. Vaccine 29:7818–25.

48. Klouwens MJ, Trentelman JJ, Ersoz JI, Nieves Marques Porto F, Sima R, Hajdusek O, Thakur M, Pal U, Hovius JW. 2021. Investigating BB0405 as a novel Borrelia afzelii vaccination candidate in Lyme borreliosis. Sci Rep 11:4775.

49. Marcinkiewicz AL, Lieknina I, Yang X, Lederman PL, Hart TM, Yates J, Chen WH, Bottazzi ME, Mantis NJ, Kraiczy P, Pal U, Tars K, Lin YP. 2020. The Factor H-Binding Site of CspZ as a Protective Target against Multistrain, Tick-Transmitted Lyme Disease. Infect Immun 88.

50. Singh P, Verma D, Backstedt BT, Kaur S, Kumar M, Smith AA, Sharma K, Yang X, Azevedo JF, Gomes-Solecki M, Buyuktanir O, Pal U. 2017. Borrelia burgdorferi BBI39 Paralogs, Targets of Protective Immunity, Reduce Pathogen Persistence Either in Hosts or in the Vector. J Infect Dis 215:1000–1009.

51. Kung F, Kaur S, Smith AA, Yang X, Wilder CN, Sharma K, Buyuktanir O, Pal U. 2016. A Borrelia burgdorferi Surface-Exposed Transmembrane Protein Lacking Detectable Immune Responses Supports Pathogen Persistence and Constitutes a Vaccine Target. J Infect Dis 213:1786–95.

52. Plotkin SA. 1993. Vaccination in the 21st century. J Infect Dis 168:29–37.

53. Bista S, Singh P, Bernard Q, Yang X, Hart T, Lin YP, Kitsou C, Singh Rana V, Zhang F, Linhardt RJ, Zhnag K, Akins DR, Hritzo L, Kim Y, Grab DJ, Dumler JS, Pal U. 2020. A Novel Laminin-Binding Protein Mediates Microbial-Endothelial Cell Interactions and Facilitates Dissemination of Lyme Disease Pathogens. J Infect Dis 221:1438–1447.

54. Kumar M, Kaur S, Kariu T, Yang X, Bossis I, Anderson JF, Pal U. 2011. Borrelia burgdorferi BBA52 is a potential target for transmission blocking Lyme disease vaccine. Vaccine 29:9012–9.

55. Sanjuan E, Esteve-Gassent MD, Maruskova M, Seshu J. 2009. Overexpression of CsrA (BB0184) alters the morphology and antigen profiles of Borrelia burgdorferi. Infect Immun 77:5149–62.

56. Caimano MJ, Iyer R, Eggers CH, Gonzalez C, Morton EA, Gilbert MA, Schwartz I, Radolf JD. 2007. Analysis of the RpoS regulon in Borrelia burgdorferi in response to mammalian host signals provides insight into RpoS function during the enzootic cycle. Mol Microbiol 65:1193–217.

57. Alanazi F, Raghunandanan S, Priya R, Yang XF. 2023. The Rrp2-RpoN-RpoS pathway plays an important role in the blood-brain barrier transmigration of the Lyme disease pathogen. Infect Immun 91:e0022723.

58. Grassmann AA, Tokarz R, Golino C, McLain MA, Groshong AM, Radolf JD, Caimano MJ. 2023. BosR and PlzA reciprocally regulate RpoS function to sustain Borrelia burgdorferi in ticks and mammals. The Journal of Clinical Investigation 133.

59. Brandt ME, Riley BS, Radolf JD, Norgard MV. 1990. Immunogenic integral membrane proteins of Borrelia burgdorferi are lipoproteins. Infect Immun 58:983–91.

60. Pal U, Kitsou C, Drecktrah D, Yaş Ö B, Fikrig E. 2021. Interactions Between Ticks and Lyme Disease Spirochetes. Curr Issues Mol Biol 42:113–144.

61. Fikrig E, Telford SR, 3rd, Barthold SW, Kantor FS, Spielman A, Flavell RA. 1992. Elimination of Borrelia burgdorferi from vector ticks feeding on OspA-immunized mice. Proc Natl Acad Sci U S A 89:5418–21.

62. Bockenstedt LK, Gonzalez DG, Haberman AM, Belperron AA. 2012. Spirochete antigens persist near cartilage after murine Lyme borreliosis therapy. The Journal of Clinical Investigation 122:2652–2660.

63. Norris SJ. 2014. vls Antigenic Variation Systems of Lyme Disease Borrelia: Eluding Host Immunity through both Random, Segmental Gene Conversion and Framework Heterogeneity. Microbiol Spectr 2.

64. Tracy KE, Baumgarth N. 2017. Borrelia burgdorferi Manipulates Innate and Adaptive Immunity to Establish Persistence in Rodent Reservoir Hosts. Front Immunol 8:116.

65. Anderson C, Brissette CA. 2021. The Brilliance of Borrelia: Mechanisms of Host Immune Evasion by Lyme Disease-Causing Spirochetes. Pathogens 10.

66. Wang Y, Kern A, Boatright NK, Schiller ZA, Sadowski A, Ejemel M, Souders CA, Reimann KA, Hu L, Thomas WD, Jr., Klempner MS. 2016. Pre-exposure Prophylaxis With OspA-Specific Human Monoclonal Antibodies Protects Mice Against Tick Transmission of Lyme Disease Spirochetes. J Infect Dis 214:205–11.

67. Zhi H, Xie J, Skare JT. 2018. The Classical Complement Pathway Is Required to Control Borrelia burgdorferi Levels During Experimental Infection. Front Immunol 9:959.

68. Phillip K, Nair N, Samanta K, Azevedo JF, Brown GD, Petersen CA, Gomes-Solecki M. 2021. Maternal transfer of neutralizing antibodies to B. burgdorferi OspA after oral vaccination of the rodent reservoir. Vaccine 39:4320–4327.

69. Azevedo JF, Joyner G, Kundu S, Samanta K, Gomes-Solecki M. 2025. Maternal transfer of oral vaccine induced anti-OspA antibodies protects Peromyscus spp. from tick-transmitted Borrelia burgdorferi. Infection and Immunity 93:e00216–25.

70. Earnhart CG, Buckles EL, Marconi RT. 2007. Development of an OspC-based tetravalent, recombinant, chimeric vaccinogen that elicits bactericidal antibody against diverse Lyme disease spirochete strains. Vaccine 25:466–480.

71. Alexopoulou L, Thomas V, Schnare M, Lobet Y, Anguita J, Schoen RT, Medzhitov R, Fikrig E, Flavell RA. 2002. Hyporesponsiveness to vaccination with Borrelia burgdorferi OspA in humans and in TLR1- and TLR2-deficient mice. Nat Med 8:878–84.

72. Bockenstedt LK, Fikrig E, Barthold SW, Flavell RA, Kantor FS. 1996. Identification of a Borrelia burgdorferi OspA T cell epitope that promotes anti-OspA IgG in mice. J Immunol 157:5496–502.

73. Luke CJ, Huebner RC, Kasmiersky V, Barbour AG. 1997. Oral delivery of purified lipoprotein OspA protects mice from systemic infection with Borrelia burgdorferi. Vaccine 15:739–46.

74. Buckles EL, Earnhart CG, Marconi RT. 2006. Analysis of antibody response in humans to the type A OspC loop 5 domain and assessment of the potential utility of the loop 5 epitope in Lyme disease vaccine development. Clin Vaccine Immunol 13:1162–5.

75. Salazar JC, Pope CD, Moore MW, Pope J, Kiely TG, Radolf JD. 2005. Lipoprotein-dependent and - independent immune responses to spirochetal infection. Clin Diagn Lab Immunol 12:949–58.

76. Earnhart CG, Buckles EL, Dumler JS, Marconi RT. 2005. Demonstration of OspC type diversity in invasive human lyme disease isolates and identification of previously uncharacterized epitopes that define the specificity of the OspC murine antibody response. Infect Immun 73:7869–77.

77. Earnhart CG, Marconi RT. 2007. An octavalent lyme disease vaccine induces antibodies that recognize all incorporated OspC type-specific sequences. Hum Vaccin 3:281–9.

78. Earnhart CG, Buckles EL, Marconi RT. 2007. Development of an OspC-based tetravalent, recombinant, chimeric vaccinogen that elicits bactericidal antibody against diverse Lyme disease spirochete strains. Vaccine 25:466–80.

79. Shoberg RJ, Thomas DD. 1993. Specific adherence of Borrelia burgdorferi extracellular vesicles to human endothelial cells in culture. Infect Immun 61:3892–900.

80. Probert WS, LeFebvre RB. 1994. Protection of C3H/HeN mice from challenge with Borrelia burgdorferi through active immunization with OspA, OspB, or OspC, but not with OspD or the 83-kilodalton antigen. Infect Immun 62:1920–6.

81. Hefty PS, Brooks CS, Jett AM, White GL, Wikel SK, Kennedy RC, Akins DR. 2002. OspE-related, OspF-related, and Elp lipoproteins are immunogenic in baboons experimentally infected with Borrelia burgdorferi and in human lyme disease patients. J Clin Microbiol 40:4256–65.

82. Dulipati V, Kotimaa J, Rezola M, Kontiainen M, Jarva H, Nyman D, Meri S. 2024. Antibody responses to immunoevasion proteins BBK32 and OspE constitute part of the serological footprint in neuroborreliosis but are insufficient to prevent the disease. Scand J Immunol 99:e13353.

83. Seshu J, Esteve-Gassent MD, Labandeira-Rey M, Kim JH, Trzeciakowski JP, Hook M, Skare JT. 2006. Inactivation of the fibronectin-binding adhesin gene bbk32 significantly attenuates the infectivity potential of Borrelia burgdorferi. Mol Microbiol 59:1591–601.

84. Promnares K, Kumar M, Shroder DY, Zhang X, Anderson JF, Pal U. 2009. Borrelia burgdorferi small lipoprotein Lp6.6 is a member of multiple protein complexes in the outer membrane and facilitates pathogen transmission from ticks to mice. Mol Microbiol 74:112–125.

85. Brooks CS, Vuppala SR, Jett AM, Akins DR. 2006. Identification of Borrelia burgdorferi outer surface proteins. Infect Immun 74:296–304.

86. Brandt KS, Patton TG, Allard AS, Caimano MJ, Radolf JD, Gilmore RD. 2014. Evaluation of the Borrelia burgdorferi BBA64 protein as a protective immunogen in mice. Clin Vaccine Immunol 21:526–33.

87. Yang X, Lenhart TR, Kariu T, Anguita J, Akins DR, Pal U. 2010. Characterization of unique regions of Borrelia burgdorferi surface-located membrane protein 1. Infect Immun 78:4477–87.

88. Li X, Neelakanta G, Liu X, Beck DS, Kantor FS, Fish D, Anderson JF, Fikrig E. 2007. Role of outer surface protein D in the Borrelia burgdorferi life cycle. Infect Immun 75:4237–44.

89. Fikrig E, Pal U, Chen M, Anderson JF, Flavell RA. 2004. OspB antibody prevents Borrelia burgdorferi colonization of Ixodes scapularis. Infect Immun 72:1755–9.

90. Anderton JM, Tokarz R, Thill CD, Kuhlow CJ, Brooks CS, Akins DR, Katona LI, Benach JL. 2004. Whole-genome DNA array analysis of the response of Borrelia burgdorferi to a bactericidal monoclonal antibody. Infect Immun 72:2035–44.

91. Katona LI, Ayalew S, Coleman JL, Benach JL. 2000. A bactericidal monoclonal antibody elicits a change in its antigen, OspB of Borrelia burgdorferi, that can be detected by limited proteolysis. J Immunol 164:1425–31.

92. Coleman JL, Rogers RC, Rosa PA, Benach JL. 1994. Variations in the ospB gene of Borrelia burgdorferi result in differences in monoclonal antibody reactivity and in production of escape variants. Infect Immun 62:303–7.

93. Probert WS, Crawford M, LeFebvre RB. 1997. Antibodies to OspB prevent infection of C3H mice challenged with Borrelia burgdorferi isolates expressing truncated OspB antigens. Vaccine 15:15–9.

94. O’Bier NS, Camire AC, Patel DT, Billingsley JS, Hodges KR, Marconi RT. 2024. Development of novel multi-protein chimeric immunogens that protect against infection with the Lyme disease agent, Borreliella burgdorferi. mBio 15:e0215924.

95. Fikrig E, Tao H, Kantor FS, Barthold SW, Flavell RA. 1993. Evasion of protective immunity by Borrelia burgdorferi by truncation of outer surface protein B. Proc Natl Acad Sci U S A 90:4092–6.

96. Li X, Strle K, Wang P, Acosta DI, McHugh GA, Sikand N, Strle F, Steere AC. 2013. Tick-specific borrelial antigens appear to be upregulated in American but not European patients with Lyme arthritis, a late manifestation of Lyme borreliosis. J Infect Dis 208:934–41.

97. Nair N, Marques A, Horn EJ, Brown G, Gomes-Solecki M. 2025. Class and isotype of VlsE-specific antibody differentiates Lyme disease stage. J Clin Microbiol 63:e0034725.

98. Beenhouwer DO, Shapiro S, Feldmesser M, Casadevall A, Scharff MD. 2001. Both Th1 and Th2 cytokines affect the ability of monoclonal antibodies to protect mice against Cryptococcus neoformans. Infect Immun 69:6445–55.

99. Belperron AA, Bockenstedt LK. 2001. Natural antibody affects survival of the spirochete Borrelia burgdorferi within feeding ticks. Infect Immun 69:6456–62.

100. Hastey CJ, Olsen KJ, Elsner RA, Mundigl S, Tran GVV, Barthold SW, Baumgarth N. 2023. Borrelia burgdorferi Infection-Induced Persistent IgM Secretion Controls Bacteremia, but Not Bacterial Dissemination or Tissue Burden. J Immunol 211:1540–1549.

101. Palmer D, Shemshadian A, Berman K, Nobles A, Willsey GG, Piazza CL, Freeman-Gallant G, Rudolph MJ, Bourgeois J, Hu L, Vance DJ, Mantis NJ. 2026. An Fc-silent OspA monoclonal antibody passively protects mice from tick and intradermal Borrelia burgdorferi challenge. PLOS ONE 21:e0339749.

102. Bhattacharya D, Mecsas J, Hu LT. 2010. Development of a vaccinia virus based reservoir-targeted vaccine against Yersinia pestis. Vaccine 28:7683–9.

103. Gingerich MC, Nair N, Azevedo JF, Samanta K, Kundu S, He B, Gomes-Solecki M. 2024. Intranasal vaccine for Lyme disease provides protection against tick transmitted Borrelia burgdorferi beyond one year. NPJ Vaccines 9:33.

104. Elias AF, Stewart PE, Grimm D, Caimano MJ, Eggers CH, Tilly K, Bono JL, Akins DR, Radolf JD, Schwan TG, Rosa P. 2002. Clonal polymorphism of Borrelia burgdorferi strain B31 MI: implications for mutagenesis in an infectious strain background. Infect Immun 70:2139–50.

105. Van Laar TA, Lin YH, Miller CL, Karna SL, Chambers JP, Seshu J. 2012. Effect of levels of acetate on the mevalonate pathway of Borrelia burgdorferi. PLoS One 7:e38171.

106. Van Laar TA, Hole C, Rajasekhar Karna SL, Miller CL, Reddick R, Wormley FL, Seshu J. 2016. Statins reduce spirochetal burden and modulate immune responses in the C3H/HeN mouse model of Lyme disease. Microbes Infect 18:430–435.

107. Chen Y, Vargas SM, Smith TC, 2nd, Karna SLR, MacMackin Ingle T, Wozniak KL, Wormley FL, Jr., Seshu J. 2021. Borrelia peptidoglycan interacting Protein (BpiP) contributes to the fitness of Borrelia burgdorferi against host-derived factors and influences virulence in mouse models of Lyme disease. PLoS Pathog 17:e1009535.

108. Smith TC, 2nd, Helm SM, Chen Y, Lin YH, Rajasekhar Karna SL, Seshu J. 2018. Borrelia Host Adaptation Protein (BadP) Is Required for the Colonization of a Mammalian Host by the Agent of Lyme Disease. Infect Immun 86.

109. Maruskova M, Seshu J. 2008. Deletion of BBA64, BBA65, and BBA66 loci does not alter the infectivity of Borrelia burgdorferi in the murine model of Lyme disease. Infect Immun 76:5274–84.

110. Maruskova M, Esteve-Gassent MD, Sexton VL, Seshu J. 2008. Role of the BBA64 locus of Borrelia burgdorferi in early stages of infectivity in a murine model of Lyme disease. Infect Immun 76:391–402.

111. Esteve-Gassent MD, Elliott NL, Seshu J. 2009. sodA is essential for virulence of Borrelia burgdorferi in the murine model of Lyme disease. Mol Microbiol 71:594–612.

112. Esteve-Gassent MD, Smith TC, 2nd, Small CM, Thomas DP, Seshu J. 2015. Absence of sodA Increases the Levels of Oxidation of Key Metabolic Determinants of Borrelia burgdorferi. PLoS One 10:e0136707.

113. Skare JT, Shang ES, Foley DM, Blanco DR, Champion CI, Mirzabekov T, Sokolov Y, Kagan BL, Miller JN, Lovett MA. 1995. Virulent strain associated outer membrane proteins of Borrelia burgdorferi. J Clin Invest 96:2380–92.

