## Supplementary material for "Protective efficacy of mutant strains of *Borrelia burgdorferi* as potential reservoir host-targeted biologics against Lyme disease": Supplementary file.docx

Supplementary Table 1: Clinical samples and the corresponding ground truth data from the Lyme Disease biobank used in this study.

| **Sample ID** | ***Days of EM Prior to Enrollment*** | ***First-tier tests*** | | | | | | ***Second-tier tests*** | | | | | | ***Two-tier Positive on 1st Draw*** |
| --- | --- | --- | --- | --- | --- | --- | --- | --- | --- | --- | --- | --- | --- | --- |
|  |  | ***ELISA*** | | ***IMMUNO BLOT*** | | | | ***ELISA*** | | ***IMMUNO BLOT*** | | | |  |
|  |  | ***Whole Cell Lysate*** | ***C-6 Peptide*** | ***IgM*** | ***IgM Bands*** | ***IgG*** | ***IgG Bands*** | ***Whole Cell Lysate*** | ***C-6 Peptide*** | ***IgM*** | ***IgM Bands*** | ***IgG*** | ***IgG Bands*** |  |
| LD515 | NA | NEG | NEG | NEG | None | NEG | p93, p41 | NA | NA | NA | NA | NA | NA | NO |
| LD526 | NA | NEG | NEG | NEG | p41 | NEG | None | NA | NA | NA | NA | NA | NA | NO |
| LD538 | NA | NEG | NEG | NEG | p41 | NEG | p58, p41 | NA | NA | NA | NA | NA | NA | NO |
| **LD585** | **10** | **POS** | **POS** | **POS** | **p41, p39, p23** | **NEG** | **p41, p39, p23** | **NA** | **NA** | **NA** | **NA** | **NA** | **NA** | **YES** |
| LD610 | NA | NEG | NEG | NEG | p41 | NEG | None | NA | NA | NA | NA | NA | NA | NO |
| LD611 | NA | NEG | NEG | NEG | p41 | NEG | p66 | NA | NA | NA | NA | NA | NA | NO |
| **LD640** | **21** | **POS** | **POS** | **NEG** | **None** | **POS** | **p93, p66, p45, p41, p39, p28** | **POS** | **POS** | **NEG** | **p41** | **POS** | **p93, p66, p45, p41, p39** | **YES** |
| **LD663** | **2** | **EQV** | **POS** | **POS** | **p41, p23** | **NEG** | **p41** | **POS** | **POS** | **POS** | **p41, p23** | **NEG** | **p41, p23** | **YES** |
| LD664 | NA | NEG | NEG | NEG | None | NEG | p93 | NA | NA | NA | NA | NA | NA | NO |
| **LD673** | **14** | **POS** | **POS** | **POS** | **p41, p23** | **POS** | **p66, p58, p41, p39, p23, p18** | **POS** | **POS** | **POS** | **p41, p23** | **NEG** | **p41, p23, p18** | **YES** |
| LD674 | NA | NEG | NEG | NEG | p41 | NEG | p58, p41 | NA | NA | NA | NA | NA | NA | NO |
| **LD677** | **1** | **POS** | **POS** | **NEG** | **p41** | **POS** | **p93, p66, p45, p41, p39, p28, p18** | **NA** | **NA** | **NA** | **NA** | **NA** | **NA** | **YES** |

Supplementary Table 2: Detailed patient information of human serum samples from the Lyme Disease biobank used in this study.

| **Sample ID** | ***Classification*** | **Age** | **Gender** | **Race** | ***EM > 5 cm at Enrollment*** | ***Days of EM Prior to Enrollment*** | ***Convalescent Draw*** | ***Whole Cell Lysate ELISA*** | ***C-6 Peptide ELISA*** | ***IMMUNO BLOT IgM*** | ***IgM Bands*** | ***IMMUNO BLOT IgG*** | ***IgG Bands*** | ***Convalescent Whole Cell Lysate ELISA*** | ***Convalescent C-6 Peptide ELISA*** | ***Convalescent IMMUNO BLOT IgM*** | ***Convalescent IgM Bands*** | ***Convalescent IMMUNO BLOT IgG*** | ***Convalescent IgG Bands*** | ***Two-tier Positive on 1st Draw*** |
| --- | --- | --- | --- | --- | --- | --- | --- | --- | --- | --- | --- | --- | --- | --- | --- | --- | --- | --- | --- | --- |
| LD515 | Control (- Serology) | 27 | Male | Hispanic or Latino | NA | NA | NA | NEG | NEG | NEG | None | NEG | p93,p41 | NA | NA | NA | NA | NA | NA | NO |
| LD526 | Control (- Serology) | 59 | Male | White | NA | NA | NA | NEG | NEG | NEG | p41 | NEG | None | NA | NA | NA | NA | NA | NA | NO |
| LD538 | Control (- Serology) | 49 | Female | Hispanic or Latino | NA | NA | NA | NEG | NEG | NEG | p41 | NEG | p58,p41 | NA | NA | NA | NA | NA | NA | NO |
| LD585 | Confirmed Lyme | 64 | Male | White | YES | 10 | NO | POS | POS | POS | p41,p39,p23 | NEG | p41,p39,p23 | NA | NA | NA | NA | NA | NA | YES |
| LD610 | Control (- Serology) | 65 | Male | White | NA | NA | NA | NEG | NEG | NEG | p41 | NEG | None | NA | NA | NA | NA | NA | NA | NO |
| LD611 | Control (- Serology) | 53 | Female | White | NA | NA | NA | NEG | NEG | NEG | p41 | NEG | p66 | NA | NA | NA | NA | NA | NA | NO |
| LD640 | Confirmed Lyme | 49 | Female | White | YES | 21 | YES | POS | POS | NEG | None | POS | p93,p66,p45,p41,p39,p28 | POS | POS | NEG | p41 | POS | p93,p66,p45,p41,p39 | YES |
| LD663 | Confirmed Lyme | 43 | Male | Hispanic or Latino | YES | 2 | YES | EQV | POS | POS | p41,p23 | NEG | p41 | POS | POS | POS | p41,p23 | NEG | p41,p23 | YES |
| LD664 | Control (- Serology) | 36 | Female | Hispanic or Latino | NA | NA | NA | NEG | NEG | NEG | None | NEG | p93 | NA | NA | NA | NA | NA | NA | NO |
| LD673 | Confirmed Lyme | 65 | Male | White | YES | 14 | YES | POS | POS | POS | p41,p23 | POS | p66,p58,p41,p39,p23,p18 | POS | POS | POS | p41,p23 | NEG | p41,p23,p18 | YES |
| LD674 | Control (- Serology) | 36 | Male | White | NA | NA | NA | NEG | NEG | NEG | p41 | NEG | p58,p41 | NA | NA | NA | NA | NA | NA | NO |
| LD677 | Confirmed Lyme | 36 | Female | Hispanic or Latino | YES | 1 | NO | POS | POS | NEG | p41 | POS | p93,p66,p45,p41,p39,p28,p18 | NA | NA | NA | NA | NA | NA | YES |

Supplementary Figure 1

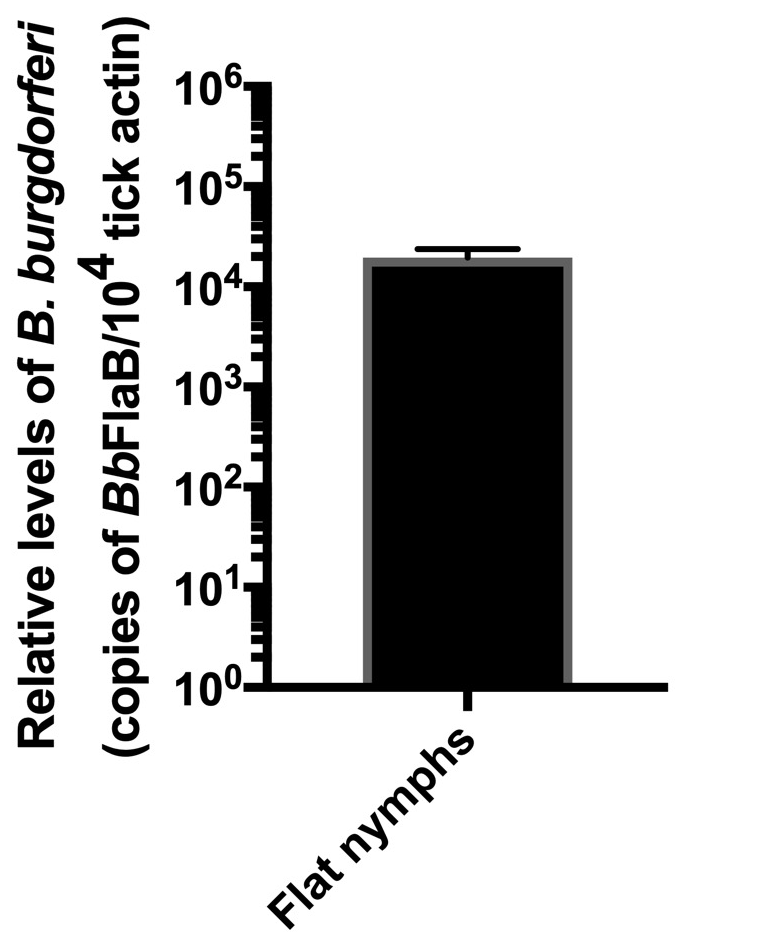

**Supplementary Figure 1: *B. burgdorferi* load in flat nymphs.** Larval ticks were allowed to feed on C3H/HeN mice that had been needle-infected with 1 × 10⁵ *B. burgdorferi* and maintained for 28 days post-infection. Engorged larvae were then allowed to molt into nymphs over a period of 6 weeks. These molted, unfed (flat) nymphs were collected for *B. burgdorferi* quantification. Genomic DNA was extracted from individual nymphs, and qPCR was performed using *BbFlaB* and tick *actin (ActB)* as targets. The bar graph shows the relative genomic load of *B. burgdorferi* in flat nymphs (n = 5), normalized to tick actin. The y-axis represents log-transformed *flaB* levels normalized to 10⁴ copies of tick actin. Error bar represents the SD demonstrating consistency in the *Bb* burden in each tested flat nymph.
